# WTAP, transcriptionally regulated by p65, promotes inflammation through m^6^A modification and phase separation

**DOI:** 10.1101/2023.10.30.564747

**Authors:** Yong Ge, Rong Chen, Tao Ling, Biaodi Liu, Jingrong Huang, Youxiang Cheng, Yi Lin, Hongxuan Chen, Xiongmei Xie, Guomeng Xia, Guanzheng Luo, Shaochun Yuan, Anlong Xu

## Abstract

Emerging evidence has linked dysregulation of *N*^6^-methyladenosine (m^6^A) to inflammation and inflammatory diseases, but the underlying mechanism still needs investigation. Here, we found that high m^6^A modification in a variety of hyperinflammatory states is p65-dependent, because Wilms tumor 1 associated protein (WTAP), a key component of the writer complex, is transcriptionally regulated by p65 and its overexpression can lead to higher m^6^A modification. Mechanistically, upregulated WTAP is more prone to phase separation to facilitate the aggregation of “writer” complex to nuclear speckles and the deposition of m^6^A marks onto transcriptionally active inflammatory transcripts, thereby accelerating proinflammatory response. Furthermore, myeloid deficiency of WTAP attenuates the severity of LPS-induced sepsis and DSS-induced IBD. Thus, the proinflammatory effect of WTAP is a general risk-increasing mechanism, and interrupting the assembly of m^6^A writer complex by targeting the phase separation of WTAP to reduce the global m^6^A level may be a potential and promising therapeutic strategy for alleviating hyperinflammation.

## Introduction

Inflammation is usually a physiological healing response that is triggered by noxious stimuli and conditions, such as infection and tissue injury (1). Moderate inflammation is essential for the pathogen clearance, tissue repair and regeneration. However, dysregulated inflammation disrupts immune homeostasis, which may lead to the development of a variety of inflammatory diseases, such as chronic inflammation, metabolic disorders, autoimmune diseases (ADs) and cancer (2, 3). To maintain immune homeostasis and prevent harmful outcomes, the activation or induction of proinflammatory regulators and effectors must be precisely controlled at multiple levels, including the transcriptional (4), posttranscriptional (5) and posttranslational (6) levels, and these are thus potential targets for the treatment of inflammatory diseases (7, 8). The therapeutic strategies for inflammatory diseases have been established and even become invaluable by developing antagonists or inhibitors against principal inflammatory effectors (NF-κB, STAT3 and JAK) and multifunctional proinflammatory cytokines (IL-6, TNF-α, IL-17A and IL-23) (9, 10), such as specific antibodies against IL-6(R) (e.g., tocilizumab and siltuximab) and TNF-α (e.g., infliximab, etanercept and adalimumab), which have been used in the treatment of many autoimmune diseases (11). However, due to pleiotropy, the therapeutic effects of targeting specific cytokines such as IL-6 or IL-17A greatly vary among different inflammatory diseases (12). In contrast, small-molecule inhibitors targeting the NF-κB and STAT3 signalling pathways lead to significant side effects that alter the homeostasis of the immune system as well as the nonimmune cells (13, 14). Therefore, in-depth research into how inflammation is precisely regulated remains particularly important for the development of new treatments against excessive inflammatory responses.

In the past two decades, studies on immune regulation have mainly focused on posttranslational modifications of proteins, such as phosphorylation and ubiquitination (6). Recently, emerging evidence has strongly indicated that immune signalling can trigger dynamic alterations to the epitranscriptome, which orchestrates the regulation of immune response (15, 16). Among those dynamic epitranscriptomics alterations, m^6^A modification is the most abundant and reversible RNA modification and can influence pre-mRNA splicing, stability, translation, location and transport (17). The m^6^A modification is mediated by a methyltransferase complex called “writer” and removed by demethylase, namely “eraser”. The writer complex composed of methyltransferase-like 3 (METTL3), methyltransferase-like 14 (METTL14), and the co-enzyme factors Wilms tumor 1 associated protein (WTAP), VIRMA (KIAA1429), ZC3H13 and RBM15, while eraser proteins include fat mass and obesity-associated protein (FTO) and alkB homolog 5 (ALKBH5) (18). Among them, WTAP may function as a regulatory subunit that binds to METTL3/14 and is required for substrate recruitment and METTL3/14 localization (19). In addition to playing roles in distinct biological processes such as embryonic development, hematopoiesis and cancer, the roles of m^6^A modification in inflammatory regulation have recently attracted intense attention (20, 21). Principal inflammatory signaling pathways, such as NF-κB, JAK-STAT and MAPK, have been found to be extensively regulated by m^6^A modification, while m^6^A related proteins have different or even opposite regulatory effects on inflammatory responses by targeting different genes or depending on specific cell and disease statuses (22, 23). For example, depleting METTL3, the key methyltransferase in the “writer” complex, reduces the m^6^A abundance on *TRAF6* mRNA and decreases the expression of TRAF6 by entrapping the transcripts in the nucleus, which inhibits the activation of the NF-κB and mitogen-activated protein kinase (MAPK) signalling pathways (24). In contrast, another study revealed that METTL3 is significantly elevated in patients with rheumatoid arthritis (RA) and that high expression of METTL3 attenuates the inflammatory response in macrophages which are stimulated with LPS (25). Furthermore, knocking-down the demethylase FTO leads to an increased overall m^6^A abundance, STAT3 phosphorylation and secretion of proinflammatory cytokines (26), whereas the knockdown of ALKBH5 inhibits cell viability and production of inflammatory cytokines (27). In addition to the m^6^A writers and erasers, m^6^A readers, such as YTHDF2, also play different or even opposite roles in regulating different inflammatory events. For example, a previous study showed that YTHDF2 limits the decay of *KDM6B* mRNA in a m^6^A-dependent manner and promotes the demethylation of histone 3 lysine 27 trimethylation (H3K27me3), which is required for the transcription of certain proinflammatory cytokines (28). YTHDF2 also accelerates the decay of m^6^A-modified transcripts encoding NF-κB-negative regulators to regulate NF-κB signaling in intratumoral Treg cells (29). However, another study indicated that YTHDF2 is an inflammatory suppressor that downregulates m^6^A-modified transcripts in inflammatory response and thereby protects hematopoietic stem cells from excessive proinflammatory signals (30). Thus, although notable evidence clearly points m^6^A modification to modulating inflammatory responses, its effects appear to depend greatly on the disease status or specific target genes.

Recently, due to the finding of aberrant RNA methylation in inflammatory diseases (21) and cancers (31), small-molecule inhibitors, which target specific “writer” or “eraser” to reprogram the m^6^A epitranscriptome, have attracted considerable attention and have been proven to be feasible (32). Notably, a research group identified a selective METTL3 catalytic inhibitor STM2457, that can suppress the growth of acute myeloid leukemia (AML) (33). Moreover, two FTO inhibitors (FB23 and FB23-2) have been designed to bind FTO directly and inhibit FTO demethylase activity, and exert a selective inhibitory effect on the proliferation of AML cells (34). In addition to the aforementioned inhibitors used to treat AML, potential inhibitors of m^6^A methylase and demethylase remain in the research stage for the treatment of inflammatory diseases. Since m^6^A methylation contributes to pluripotency, identification of the intrinsic principles underlying the mechanism through which m^6^A regulates inflammation and inflammatory diseases is extremely important before a therapeutic approach can be developed to target the m^6^A mark. The investigation of other general regulators, in addition to m^6^A methylase and demethylase inhibitors, may provide another method for developing effective and even safer treatments for inflammatory diseases.

By revealing the basic principle of m^6^A modification in inflammatory regulation, we report a new function of the m^6^A “writer” protein WTAP in controlling inflammatory responses and associated diseases. We found that WTAP is an NF-κB p65-stimulated gene and that its expression is significantly upregulated in response to a variety of inflammatory stimuli and in many inflammatory diseases. Mechanistically, after an increase in its concentration, WTAP spontaneously undergoes phase separation, which facilitates the aggregation of the “writer” complex to nuclear speckles and the deposition of m^6^A onto transcriptionally active proinflammatory genes. Hence, WTAP enhances the protein synthesis of many m^6^A-modified proinflammatory transcripts, including IL6ST, IL18R1, IL15RA, IL-1A, IL12B and CXCL11, in response to inflammatory stimuli, which accelerates the process of inflammatory responses and aggravates the severity of many inflammatory diseases, such as LPS-induced sepsis, IBD and psoriasis. Thus, our study here identified WTAP as a risk factor in inflammatory responses and thereby provides mechanistic insights into WTAP as a novel and potential therapeutic target for preventing excessive inflammation.

## Results

### Hyperinflammation is accompanied by elevated WTAP levels in many inflammatory diseases

To reveal how m^6^A modification plays an essential role in inflammation and inflammatory diseases, we analyzed data obtained from the Gene Expression Omnibus (GEO) dataset (GSE13887/137268/69063/97779/166388/208303, Supplementary Table 1), and found that among m^6^A related proteins, the expression of WTAP is commonly upregulated in patients with SLE, asthma, sepsis, rheumatoid arthritis, psoriasis, or Crohn’s disease (Supplementary Figure 1, A-F). To verify this phenomenon, we then performed statistical analyses of previously published microarray datasets (GSE19315/198326/2411/2638/227851/189847, Supplementary Table 1) and further identified that only the mRNA abundance of *WTAP* among the members of the writer complex was significantly increased after LPS stimulation in THP-1 cells (Supplementary Figure 1G) and human macrophages (Supplementary Figure 1H). Similar results were obtained from the lung tissue of mice with LPS-induced lung injury (Supplementary Figure 1I). In addition, we found that the mRNA abundance of *WTAP* was also increased in TNF-α stimulated human microvascular endothelial cells (HMECs) (Supplementary Figure 1J), *Mycobacterium tuberculosis* (H37Rv)-infected THP-1 cells (Supplementary Figure 1K) and *Salmonella typhimurium* (SL1344)-infected macrophages (Supplementary Figure 1L). These observations suggest that the overexpression of WTAP is a common phenomenon in hyperinflammation states.

To verify the abovementioned observations, THP-1 cells were treated with the Toll-like receptor 4 (TLR4) agonist LPS, an ideal reagent to activate the inflammatory signalling cascade in vivo (35). RNA-sequencing (RNA-seq) analysis was then performed and the results confirmed that only the mRNA abundance of *WTAP* among the members of the writer complex was significantly upregulated in the LPS-stimulated THP-1 cells (Figure 1A). Further qRT‒PCR (Figure 1, B-D) or immunoblotting (Figure 1, E-G) analyses conducted with THP-1 cells, peripheral blood mononuclear cells (PBMCs) and mouse bone marrow-derived macrophages (BMDMs) after LPS stimulation confirmed that the expression of WTAP was upregulated. To determine whether the upregulated expression of WTAP is ubiquitous upon specific inflammatory stress, we treated THP-1 cells and BMDMs with different TLR agonists or heat-killed bacteria, such as CL097 (a TLR7/8 ligand), Pam3CSK4 (a TLR1/2 ligand), heat-killed *Salmonella typhimurium* (HKST) and heat-killed *Listeria monocytogenes* (HKLM), and found that the mRNA and protein abundance of WTAP were both significantly increased (Figure 1, H-M). Thus, high WTAP expression is positively correlated with hyperinflammation states, implying a potential role of WTAP in the regulation of inflammation.

**Figure 1.**
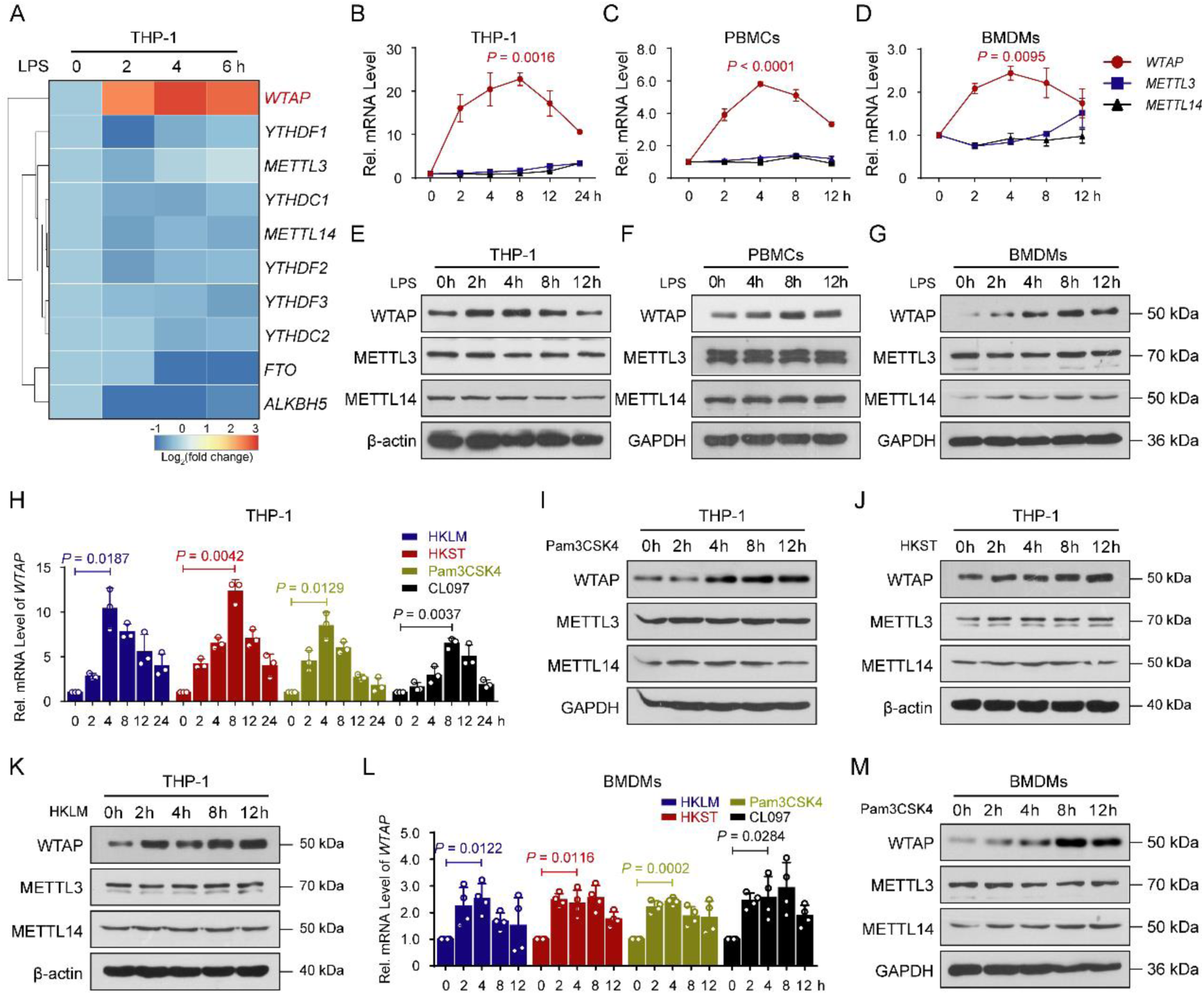
Multiple inflammatory stimuli markedly increased the expression of WTAP in macrophages. (**A**) Heatmap showing a change in the mRNA abundance of m^6^A-related genes in THP-1 cells stimulated with lipopolysaccharide (LPS) at the indicated time points. (**B** to **D**) qRT-PCR showing the mRNA abundance of the METTL3/METTL14/WTAP heterotrimer in THP-1 cells (B), peripheral blood mononuclear cells (PBMCs) (C) and bone marrow-derived macrophages (BMDMs) (D) stimulated with LPS at the indicated time points. (**E** to **G**) Immunoblots showing the protein abundance of the METTL3/METTL14/WTAP heterotrimer in THP-1 cells (E), PBMCs (F) and BMDMs (G) stimulated with LPS at the indicated time points. (H) qRT-PCR showing the mRNA abundance of WTAP in THP-1 cells stimulated with CL097, Pam3CSK4, heat-killed *Salmonella typhimurium* (HKST) or heat-killed *Listeria monocytogenes* (HKLM) at the indicated time points. (**I** to **K**) Immunoblots showing the protein abundance of the METTL3/METTL14/WTAP heterotrimer in THP-1 cells stimulated with Pam3CSK4 (I), HKST (J) or HKLM (K) at the indicated time points. (**L**) qRT-PCR showing the mRNA abundance of WTAP in BMDMs stimulated with CL097, Pam3CSK4, HKST or HKLM at the indicated time points. (**M**) Immunoblots showing the protein abundance of the METTL3/METTL14/WTAP heterotrimer in BMDMs stimulated with Pam3CSK4 at the indicated time points. Data are presented as the mean ± s.d. in (B to D), (H) and (L), with individual measurements overlaid as dots, statistical analysis was performed using a two-tailed Student’s *t*-test. Data are representative of three independent biological experiments in (E to G), (I to K) and (M).

### Upregulation of WTAP in hyperinflammation is controlled by NF-κB p65

Protein abundance can be regulated at the transcriptional, translational, or posttranslational level. To reveal the specific mechanism underlying the upregulation of WTAP at both the mRNA and protein levels upon inflammatory stimulation, bioinformatics analyses using the Promoter 2.0 prediction server (http://www.cbs.dtu.dk/services/promoter/), CpG plot (http://www.ebi.ac.uk/Tools/seqstats/emboss_cpgplot/) and JASPAR (http://jaspar.genereg.net/) were performed to identify the region between −800 and +250 in the genomic sequence of human *WTAP* containing the TATA box, CAAT box and GC box, which are characteristic of promoters. Further analyses of transcription factor-binding sites revealed that the *WTAP* promoter contains NF-κB p65-, C/EBPβ-, IRF3-, STAT3- and HIF1α-binding motifs (Supplementary Table 2). Studies have reported that both STAT3 and HIF1α can transcriptionally upregulate the expression of WTAP in some cancer cells (36, 37), but we found that LPS-induced upregulation of WTAP was not affected by SC144 treatment, which inhibited the activation of STAT3 signaling by binding IL6ST (38) (Supplementary Figure 2, A and B). Similarly, the accumulation of HIF1α induced by CoCl_2_ can upregulate WTAP (Supplementary Figure 2C), but inflammatory stimuli did not cause the accumulation of HIF1α (Supplementary Figure 2D). These data indicated that STAT3 and HIF1α have little effect on the transcriptional upregulation of WTAP under inflammatory stress. So, we next explored the transcriptional activation of WTAP by NF-κB p65, C/EBPβ and IRF3, and the p65-binding motifs exhibit the highest prevalence and are distributed in a relatively concentrated and overlapping region among these binding motifs (Figure 2A). Similar results were obtained from an analysis of the mouse *Wtap* promoter region (Figure 2B). As shown in Figure 2A and 2B, we constructed a series of reporter plasmids containing the wide-type (WT) WTAP promoter with NF-κB p65-, C/EBPβ-, IRF3-binding motifs or mutated or deleted p65-binding motifs based on the pGL3 basic construct. These reporter plasmids were then cotransfected with increasing amounts of NF-κB p65-, C/EBPβ- or IRF3-expressing plasmids into 293T cells. The results revealed that NF-κB p65 but not IRF3, increased the expression of the respective *WTAP* promoter reporter in a dose-dependent manner (Figure 2C; and Supplementary Figure 2, E and F). However, deletion or mutation of the core p65-binding sites in the *WTAP* promoter inhibited the expression of the reporter gene (Figure 2D). Similar results were obtained using reporter plasmids that contained the mouse *Wtap* promoter (Supplementary Figure 2G). Furthermore, the treatment of THP-1 cells or PBMCs with the p65 inhibitor PG490 (triptolide) inhibited the induction of WTAP by LPS (Figure 2, E-H). Consistently, using SN50, inhibitor of NF-κB p65 translocation (39), obtained the same results (Supplementary Figure 2, H and I). Next, we generated *RELA*^-/-^ (encoding p65) THP-1 cells using the CRISPR-mediated genome editing approach (Supplementary Figure 2J), and found that the upregulation of WTAP was inhibited in *RELA*^-/-^ THP-1 cells after LPS, Pam3CSK4, HKST or HKLM stimulation (Figure 2, I and J and Supplementary Figure 2, K-N). Using primers targeting p65-binding sites in the WTAP promoter for ChIP-qPCR, we found a significant enrichment of p65-binding sequence immunoprecipitated by p65 antibody compared with negative control IgG after LPS treatment (Figure 2, K and L and Supplementary Figure 2O). DNA pull-down assays also confirmed the direct binding of Flag-tagged p65 to biotin-labelled p65-binding probes of the *WTAP* promoter in vitro (Figure 2, M and N). In addition, C/EBPβ can also slightly activate the expression of WTAP (Supplementary Figure 2E), and DNA pull-down assays confirmed the direct binding of C/EBPβ to the *WTAP* promoter (Supplementary Figure 2P). Because p65 can further activate some inducible transcription factors such as ATF3, C/EBPδ and C/EBPβ, to enhance the LPS-induced transcriptional response (4), we hypothesized that although C/EBPβ may play some role in the transcription of *WTAP*, the upregulation of *WTAP* predominantly depends on the activation of NF-κB p65.

**Figure 2.**
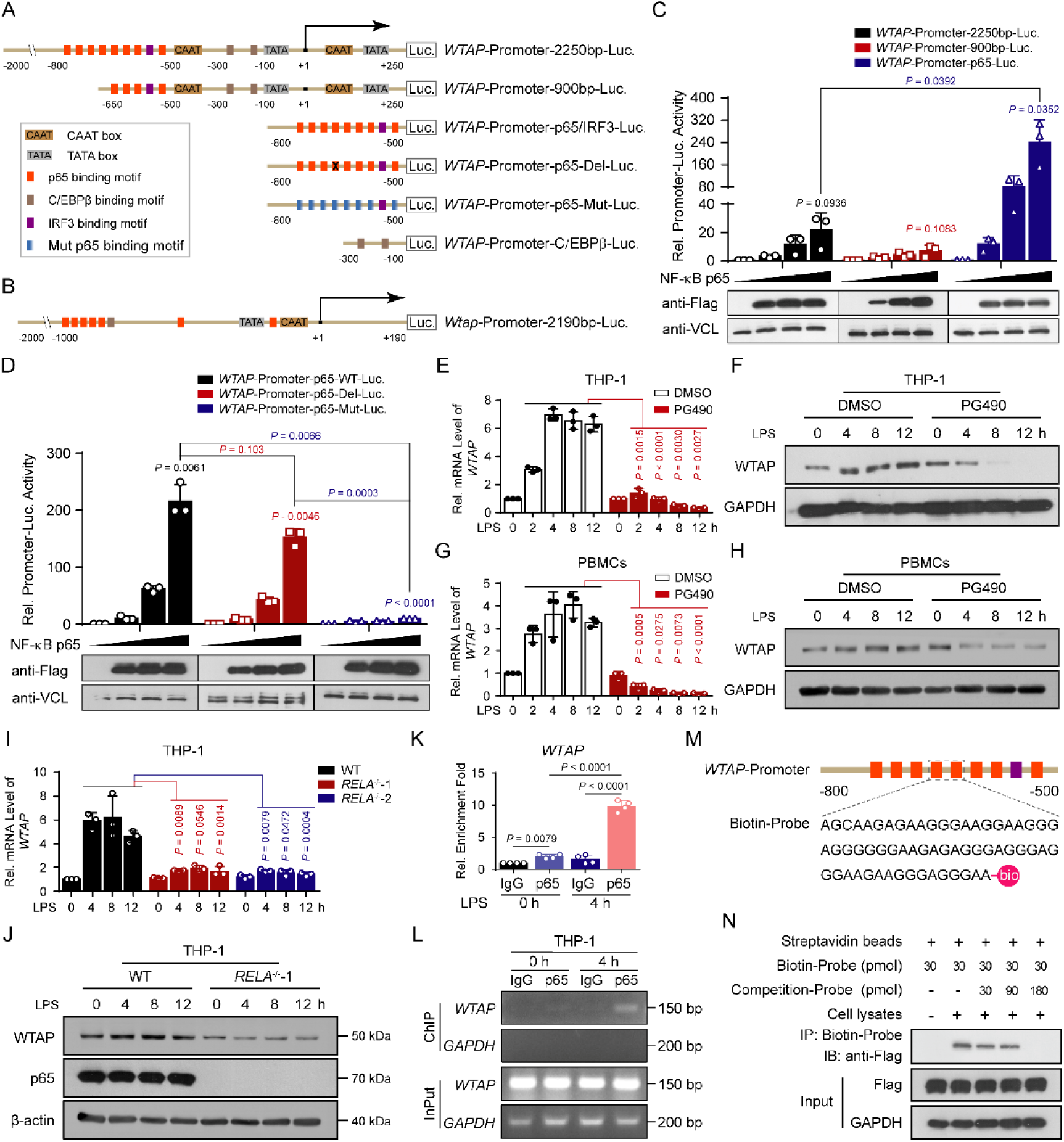
Upregulation of WTAP in hyperinflammation is controlled by NF-κB p65. (**A**) Schematic of the full-length, mutant and truncated human *WTAP* promoter. (**B**) Schematic of the mouse *Wtap* promoter. (**C** and **D**) Luciferase activity analyses in 293T cells transfected with a luciferase reporter for the *WTAP* promoter with WT (**C**), mutant or truncated (**D**) p65-binding motif, together with increasing doses of Flag-tagged NF-κB p65. (**E** to **H**) qRT-PCR or immunoblots showing the expression of WTAP in THP-1 cells (E and F) or PBMCs (G and H) pretreated with PG490, followed by stimulation with LPS at the indicated time points. (**I** and **J**) qRT-PCR (I) or immunoblots (J) showing the expression of WTAP in WT and *RELA*^-/-^ THP-1 cells stimulated with LPS at different time points. (**K** and **L**) ChIP‒qPCR (K) and Semi-quantitative RT‒PCR (L) assays analyzing p65 occupancy on the WTAP promoter in THP-1 cells before and after LPS treatment. (**M**) Schematic showing the biotin-labeled *WTAP* promoter probes. (**N**) DNA pull-down assays showing the effect of NF-κB p65 binding with *WTAP* promoter probes. Data are presented as the mean ± s.d. in (C to E), (G), (I) and (K), with individual measurements overlaid as dots, statistical analysis was performed using a two-tailed Student’s *t*-test. Data are representative of three independent biological experiments in (C and D), (F), (H), (J), (L) and (N).

### WTAP positively regulates proinflammatory responses

To determine the significance of WTAP in inflammatory responses, we first generated *WTAP*-knockout THP-1 cells using the CRISPR/Cas9 approach. Two sgRNAs were designed to target the second and third exons (E2 and E3) of *WTAP*. sgRNA#1 caused two bases insertion, whereas sgRNA#2 caused one base deletion in the *WTAP* genome locus (Supplementary Figure 3, A and B), leading to the introduction of a new stop codon upstream of the CDS and resulting in an early termination of WTAP translation. However, with open reading frame (ORF) prediction, it was found that the cells may skip these mutations by alternative translation initiation to produce alternative WTAP isoforms with 376 and 319 aa, respectively (Supplementary Figure 3C). A subsequent immunoblotting assay using a monoclonal antibody (mAb) purchased from Abcam confirmed the extremely low expression of these two isoforms (Supplementary Figure 3D). Thus, the cell lines were designated as “*WTAP^Δ1-20^* THP-1 cells” and “*WTAP^Δ1-77^* THP-1 cells”. In addition to generating KO THP-1 cells, we also used the same sgRNAs to knockout WTAP in 293T cells and successfully obtained clones using sgRNA#1. Genomic sequencing confirmed that sgRNA#1 targeting the second exon introduced four bases deletion, also resulting in an early termination of WTAP translation. But just like what happened in *WTAP^Δ1-20^* THP-1 cells, the truncated WTAP^Δ1-20^ isoform may be produced and rarely expressed in 293T knockout cells (Supplementary Figure 3, E-G), thus the cells were designated as “*WTAP^Δ1-20^* 293T cells”. We next generated the *Wtap* conditional KO (CKO) mice by crossing *Wtap*^flox/flox^ mice with mice expressing Cre recombinase under the control of the lysozyme 2 promoter (*LyzM*-Cre) (Supplementary Figure 3H). Genomic sequencing confirmed that the third exon was fully removed after crossing, which also resulted in early termination of WTAP translation (Supplementary Figure 3, I and J). Similarly, ORF prediction showed that gene-edited BMDMs may also produce alternative WTAP isoforms with 319 aa (Supplementary Figure 3J). However, none of the purchased antibodies targeting WTAP can identify this predicted isoform (Supplementary Figure 3, K and L). Although there are not enough commercially available antibodies for detection, the expression of WTAP^Δ1-77^ should be extremely low in BMDMs if present, like what we have observed in THP-1 cells.

It has been demonstrated that WTAP possesses a nuclear localization signal (NLS) at its N terminus (Supplementary Figure 3C), and mutation of this signal can affect the entry of the protein into the nucleus and its function (40). Since the NLS is missing in both alternative isoforms, their entry to the nucleus may be affected. The nuclear-cytoplasmic extraction assays confirmed the WTAP^Δ1-20^ and WTAP^Δ1-77^ were more concentrated in the cytoplasm (Supplementary Figure 3, M and N). Moreover, LC‒MS/MS assays indicated that ectopic expression of full-length WTAP but not the two alternative isoforms in *WTAP^Δ1-20^* 293T cells could increase the global m^6^A modification level (Supplementary Figure 3O). Similarly, the m^6^A modification levels of *WTAP^Δ1-20^* and *WTAP^Δ1-77^* THP-1 cells were significantly lower than those of wild-type cells (Supplementary Figure 3P), the same trend was observed in BMDMs from *LyzM*-Cre^+^ *Wtap^Δ1-77^* mice (Supplementary Figure 3Q).

Due to the extremely low expression and the severely impaired function of WTAP^Δ1-20^ and WTAP^Δ1-77^, the status of *WTAP^Δ1-20^* and *WTAP^Δ1-77^* cells should be very close to that of cells in which the protein is completely knocked out. Additionally, no differences in the proportion of major immune cell populations were observed between *Wtap*^fl/fl^ and *LyzM*-Cre^+^ *Wtap^Δ1-77^*mice in steady state (Supplementary Figure 4, A-C), indicating that the depletion of *Wtap* in myeloid cells did not affect macrophage development and maturation.

After verifying the status of WTAP KO THP-1 cells and CKO BMDMs, we next performed RNA-seq analyses with WT and *WTAP^Δ1-77^*THP-1 cells before and after LPS treatment. KEGG enrichment analyses showed that the transcripts downregulated in *WTAP^Δ1-77^* THP-1 cells were mainly enriched in the cytokine production and inflammatory signalling pathways (Figure 3, A and B). In contrast, KEGG analyses of the genes upregulated in *WTAP^Δ1-77^* THP-1 cells showed that there were enriched in a few pathways that are not directly associated with inflammatory responses (Supplementary Figure 5A). To confirm the results obtained by RNA-seq, qRT‒PCR analyses were performed and revealed marked reductions in *IL6*, *CCL2*, *CCL8* and *CXCL8* expression in *WTAP^Δ1-77^* THP-1 cells after stimulation with LPS (Figure 3 C). Similar results were obtained with *WTAP^Δ1-77^* THP-1 cells treated with Pam3CSK4, HKST or HKLM (Supplementary Figure 5B). Consistently, *WTAP^Δ1-77^* THP-1 cells exhibited a significant reduction in IL-6 expression and secretion in response to the aforementioned stimuli compared with the WT cells (Figure 3, D and E). Furthermore, ectopic expression of WTAP (Supplementary Figure 5C) substantially facilitated the expression of IL-6 and other inflammatory genes under inflammatory stimulation (Supplementary Figure 5, D and E). These results confirmed that WTAP accelerates inflammatory responses by promoting the expression of many proinflammatory cytokines under different inflammatory stimuli.

**Fig. 3.**
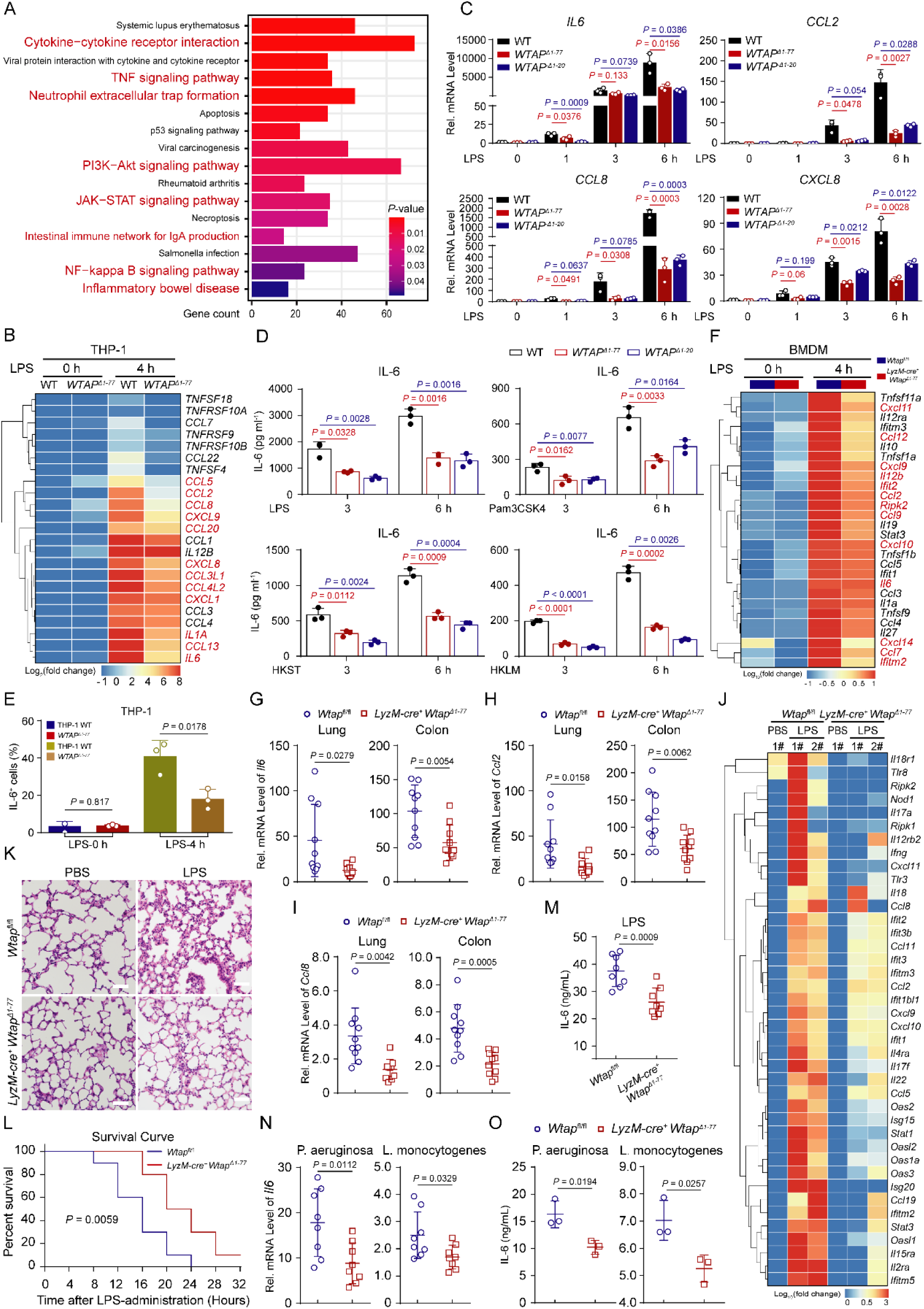
WTAP positively regulates proinflammatory responses. (**A**) KEGG analyses of down-regulated genes in *WTAP^Δ1-77^* THP-1 cells during LPS-induced inflammatory stress compared with WT cells. (**B**) Heatmap showing a change in the mRNA abundance of pro-inflammatory cytokines in WT and *WTAP^Δ1-77^* THP-1 cells stimulated with LPS at the indicated time points. (**C**) qRT–PCR showing the mRNA abundance of *IL6*, *CCL2*, *CCL8* and *CXCL8* in WT and *WTAP^Δ1-77^* THP-1 cells stimulated with LPS at the indicated time points. (**D**) ELISAs were performed to detect IL-6 secretion in supernatants of WT and *WTAP^Δ1-77^*THP-1 cells stimulated with LPS, Pam3CSK4, HKST or HKLM at 3 and 6 hr. (**E**) Representative flow cytometry data showing IL-6 fluorescence intensity in WT and *WTAP^Δ1-77^* THP-1 cells before and after LPS stimulation. (**F**) Heatmap showing a change of mRNA abundance of pro-inflammatory cytokines in WT and *Wtap^Δ1-77^*BMDMs stimulated with LPS at the indicated time points. (**G** to **I**) qRT–PCR showing the mRNA abundance of *Il6* (G), *Ccl2* (H) and *Ccl8* (I) in the lung or colon tissues from *Wtap*^fl/fl^ and *LyzM-cre*^+^ *Wtap^Δ1-77^*mice that were intraperitoneally injected with LPS (10 mg kg^-1^) for 12 hr. n = 10 mice per group. (**J**) Heatmap showing a change of mRNA abundance of pro-inflammatory cytokines in the colon tissues from *Wtap*^fl/fl^ and *LyzM-cre*^+^ *Wtap^Δ1-77^* mice that were intraperitoneally injected with LPS for 12 hr. (**K**) Hematoxylin & eosin staining (H&E) assays showing the lung injury of the *Wtap*^fl/fl^ and *LyzM-cre*^+^ *Wtap^Δ1-77^* mice that were intraperitoneally injected with LPS (10 mg kg^-1^) for 6 hr. Scale bars, 50 μm. n = 4 mice per group. (**L**) The survival of *Wtap*^fl/fl^ and *LyzM-cre*^+^ *Wtap^Δ1-77^* mice was monitored for 32 hr following the intraperitoneal injection of LPS (40 mg kg^-1^). n = 10 mice per group. (**M**) ELISAs were performed to detect IL-6 secretion in serum from *Wtap*^fl/fl^ and *LyzM-cre*^+^ *Wtap^Δ1-77^* mice that were intraperitoneally injected with LPS (10 mg kg^-1^) for 12 hr. n = 8 mice per group. (**N**) qRT–PCR showing the mRNA abundance of *Il6* in the lung tissues from *Wtap*^fl/fl^ and *LyzM-cre*^+^ *Wtap^Δ1-77^* mice that were intraperitoneally injected with *P. aeruginosa* or *L. monocytogenes* for 16 hr. n = 8 mice per group. (**O**) ELISAs were performed to detect IL-6 secretion in serum from *Wtap*^fl/fl^ and *LyzM-cre*^+^ *Wtap^Δ1-77^* mice as in that were intraperitoneally injected with *P. aeruginosa* or *L. monocytogenes* for 16 hr. n = 3 mice per group. Data are presented as the mean ± s.d. in (C to E), (G to I), and (M to O), with individual measurements overlaid as dots, statistical analysis was performed using a two-tailed Student’s *t*-test. Data are representative of three independent biological experiments in (K).

### WTAP aggravates LPS-induced sepsis in mice

Because WTAP was upregulated after LPS stimulation and WTAP deficiency can reduce inflammatory responses in macrophages, we next examined the biological effects of WTAP on the progression of inflammatory diseases in *LyzM*-Cre^+^ *Wtap^Δ1-77^*mice. BMDMs first isolated from *Wtap*^fl/fl^ and *LyzM*-Cre^+^ *Wtap^Δ1-77^* mice were treated with LPS and then used for RNA-seq analyses. The results showed that WTAP depletion in BMDMs exhibited a significant reduction in the expression of proinflammatory cytokines upon LPS stimulation (Figure 3F), which is consistent with the trend observed in THP-1 cells. As mentioned above, the *WTAP* mRNA abundance was significantly higher in the PBMCs of sepsis patients than in those of healthy ones (Supplementary Figure 1C). A retrospective study of publicly available data also revealed that WTAP is a risk gene for sepsis, its expression was significantly correlated with 28-day cumulative mortality (41). Thus, we further investigated the biological effects of WTAP using *LyzM*-Cre^+^ *Wtap^Δ1-77^* and *Wtap*^fl/fl^ mouse models of LPS-induced sepsis. Consistent with the results obtained from LPS-induced macrophages, the LPS-triggered expression of *Il6*, *Ccl2* and *Ccl8* in lung and colon tissues was decreased in *LyzM*-Cre^+^ *Wtap^Δ1-77^* mice compared with *Wtap*^fl/fl^ mice (Figure 3, G-I), whereas the expression of *Il1a*, *Il1b* and *Tnfa* showed no significant differences (Supplementary Figure 5, F and G). The same trend was obtained with RNA-seq data from colon samples (Figure 3J). Notably, *LyzM*-Cre^+^ *Wtap^Δ1-77^* mice showed less lung inflammation and fewer pathological characteristics of lung injury (Figure 3K) and were therefore more tolerant to LPS-induced fatal sepsis (Figure 3L) than their WT counterparts. Moreover, the serum concentration of IL-6 was significantly reduced in *LyzM*-Cre^+^ *Wtap^Δ1-77^*mice compared with their *Wtap*^fl/fl^ littermates (Figure 3M). Similarly, after intraperitoneal injection of *Pseudomonas aeruginosa* (*P. aeruginosa*) or *Listeria monocytogenes* (*L. monocytogenes*), the induced expression of *Il6*, *Ccl2* and *Ccl8* in lung (Figure 3N and Supplementary Figure 5, H and I) and colon (Supplementary Figure 5, J-L) tissues, as well as the secretion of IL-6 into serum (Figure 3 O), were decreased in *LyzM*-Cre^+^ *Wtap^Δ1-77^* mice. Collectively, these data suggested that the increased expression of WTAP aggravates LPS- or bacteria-induced inflammatory responses in vivo.

### WTAP promotes activation of the STAT3 signalling axis through the m^6^A modification to accelerate inflammatory responses

As a key m^6^A “writer”, WTAP is required for anchoring METTL3 and other cofactors to nuclear speckles to modulate m^6^A modification of RNA (19). To determine whether WTAP affects the inflammatory responses through m^6^A modification, we first quantified m^6^A abundance in cells by LC–MS/MS assays, and found that the overall m^6^A modification level increased in WT cells but not WTAP-deficient cells after LPS stimulation (Figure 4, A and B). These data suggested that upregulation of WTAP can significantly improve the overall m^6^A modification level during inflammatory stimuli, which is consistent with the previous conclusion that WTAP is responsible for the observed increase in m^6^A modification upon bacterial infection (28).

**Figure 4.**
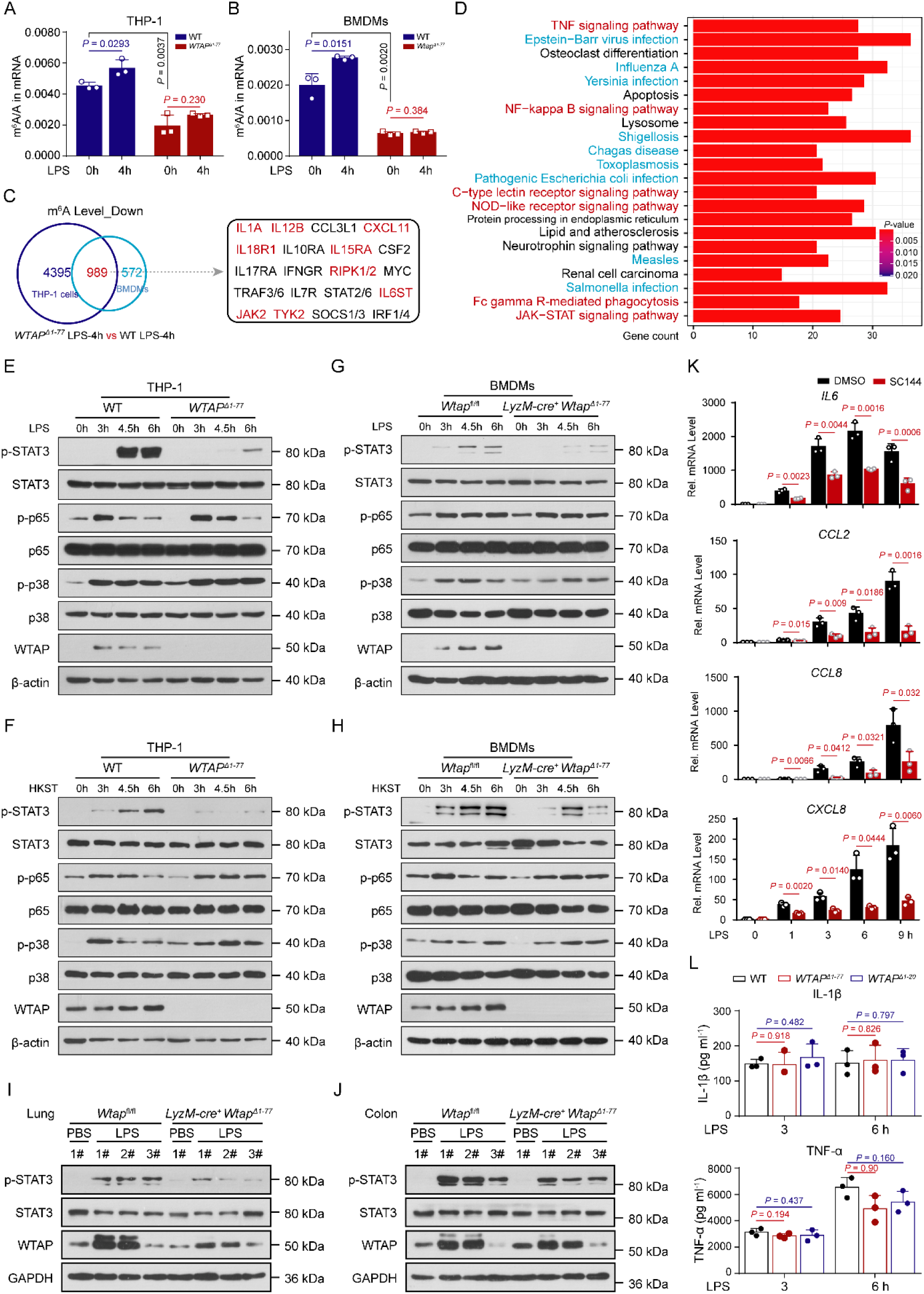
WTAP promotes the activation of STAT3 signaling through the m^6^A modification to accelerate inflammatory responses. (**A** and **B**) LC–MS/MS quantification of m^6^A abundance in mRNA extracted from WT and *WTAP^Δ1-77^* THP-1 cells or *Wtap^Δ1-77^*BMDMs with or without LPS treatment. (**C**) Venn diagrams showing transcripts with decreased m^6^A abundance in *WTAP^Δ1-77^* THP-1 cells and *Wtap^Δ1-77^*BMDMs compared with WT cells stimulated with LPS for 4 hr. (**D**) KEGG enrichment analysis of the overlapping transcripts presented in (C). (**E** to **H**) Immunoblots showing total and phosphorylated (p-) STAT3, p65 and p38 levels in WT and *WTAP^Δ1-77^* THP-1 cells (E and F) or BMDMs (G and H) stimulated with LPS or HKST at the indicated time points. (**I** and **J**) Immunoblots showing the total and phosphorylated STAT3 level in the lung (I) or colon (J) tissues from *Wtap*^fl/fl^ and *LyzM-cre*^+^ *Wtap^Δ1-77^* mice that had been intraperitoneally injected with LPS (10 mg kg^-1^) for 6 hr. (**K**) qRT–PCR showing the expression of *IL6*, *CCL2*, *CCL8* and *CXCL8* in THP-1 cells treated with SC144, followed by stimulation with LPS at the indicated time points. (**L**) ELISAs were performed to measure IL-1β and TNF-α secretion in supernatants of WT and *WTAP^Δ1-77^* THP-1 cells that were stimulated with LPS at 3 and 6 hr. Data are presented as the mean ± s.d. in (A), (B), (K) and (L), with individual measurements overlaid as dots, statistical analysis was performed using a two-tailed Student’s *t*-test. Data are representative of three independent biological experiments in (E to J).

We then performed methylated RNA immunoprecipitation sequencing (MeRIP-Seq) analyses to map mRNA transcripts with different m^6^A peaks accompanied by different mRNA levels in WT and *WTAP^Δ1-77^* THP-1 cells or *Wtap^Δ1-77^* BMDMs after LPS stimulation. According to the MeRIP-Seq data, the consensus m^6^A core motifs were enriched in the m^6^A peaks in all the samples (Supplementary Figure 6A). A peak distribution analysis showed that the m^6^A sites were enriched in both exons and 3’UTRs (Supplementary Figure 6B), with the highest enrichment near the stop codon (Supplementary Figure 6, C and D). Moreover, the MeRIP-Seq analysis revealed that the m^6^A peaks of nearly 5,000 or more than 1,500 transcripts were lower in *WTAP^Δ1-77^*THP-1 cells or in *Wtap^Δ1-77^* BMDMs compared with those in WT cells upon LPS treatment, respectively (Figure 4C). These identified transcripts were enriched in many inflammatory signalling pathways, including the NF-κB (TNF-α/TLR/NLR/CLR-mediated), JAK-STAT and MAPK pathways (Supplementary Figure 6, E and F). We further compared the transcripts with reduced m^6^A marks after WTAP deletion between THP-1 cells and BMDMs treated with LPS and identified 989 overlapping transcripts (Figure 4C). KEGG enrichment analyses revealed that these overlapping transcripts were closely related to bacterial infection and inflammatory signalling pathways (Figure 4D), such as *IL6ST*, *IL15RA*, *IL18R1*, *TYK2*, *JAK2*, *RIPK2*, *IL1A*, *IL12B* and *CXCL11* (Figure 4C).

Since both RNA-seq and MeRIP-seq indicated that WTAP is closely related to inflammatory pathways, such as the NF-κB, JAK-STAT and MAPK pathways, we then explored the effect of WTAP on the activation of these pathways. The results showed that the phosphorylation level of STAT3, but not that of p65 or p38, was significantly decreased in *WTAP^Δ1-77^* THP-1 cells in response to LPS or HKST treatment (Figure 4, E and F and Supplementary Figure 6, G and H). To further confirm this effect, the same treatments were performed with BMDMs from *Wtap^fl/fl^*or *LyzM*-Cre^+^ *Wtap^Δ1-77^* mice, and similar results were obtained (Figure 4, G and H and Supplementary Figure 6, G and H). Consistently, i.p. injection of LPS upregulated the phosphorylation level of STAT3 (Figure 4, I and J). However, such induction was significantly reduced in the lung (Figure 4I) and colon (Figure 4J) tissues from *LyzM*-Cre^+^ *Wtap^Δ1-77^*mice. On the contrary, ectopic expression of WTAP in *WTAP^Δ1-77^* THP-1 cells substantially facilitated the phosphorylation level of STAT3 after LPS stimulation (Supplementary Figure 6I). We further found that the LPS-induced mRNA expression of *IL6*, *CCL2*, *CCL8* and *CXCL8* (Figure 4K), but not that of *TNFA*, *IL1B* and *LTA* (Supplementary Figure 6J), was markedly reduced after pretreatment with gp130 (IL6ST) inhibitor SC144. Interestingly, although the expression of many proinflammatory cytokines was reduced in *WTAP^Δ1-77^* THP-1 cells, the expression and secretion of IL-1β, TNF-α or LTA (TNF-β) showed no significant change compared with those of WT THP-1 cells in response to the indicated stimuli (Figure 4L and Supplementary Figure 7, K-M), which is consistent with the fact that p65 activation is not affected by WTAP. Together, these data suggested that WTAP-mediated m^6^A modification is deeply involved in the regulation of bacterial infection and inflammatory response in both human and mice and may accelerate inflammatory responses by enhancing the STAT3 signalling axis.

### WTAP promotes the protein output of proinflammatory genes through m^6^A modification

Consistent with our experimental results, many m^6^A-modified genes that were found to be regulated by WTAP based on the sequencing data are involved in the activation and regulation of the STAT3 signalling axis, such as IL6ST, TYK2, JAK2, IL15RA and IL18R1 (Figure 4C and Supplementary Figure 7, A-L). Since the transcription and secretion of IL-6 is significantly reduced in our analyses and its contribution to the activation of STAT3 may be predominant in acute inflammatory response, we then further measured the protein abundance of the identified key components involved in the IL-6/STAT3 signalling axis, including IL6ST, IL6R, TYK2, JAK1, JAK2 and STAT3 (42). The results showed that the protein abundance of IL6ST but not the other tested components was reduced in *WTAP^Δ1-77^* THP-1 cells and *Wtap^Δ1-77^* BMDMs compared with control cells (Figure 5A and Supplementary Figure 8A). Moreover, the upregulation of IL6ST induced by LPS was inhibited in the *WTAP^Δ1-77^* THP-1 cells (Figure 5B). Consistently, the expression of IL6ST was significantly reduced on the cell surface of *Wtap^Δ1-77^*macrophages (Figure 5C). Notably, the analysis of mouse models of LPS-induced sepsis showed that the increased protein expression of IL6ST and WTAP (Supplementary Figure 8, B and C) was reduced in the lung and colon tissues (Figure 5D) from *LyzM*-Cre^+^ *Wtap^Δ1-77^* mice. We next generated *IL6ST*^-/-^ THP-1 cells using the CRISPR/Cas9 approach, and as shown in *WTAP^Δ1-77^* THP-1 cells, the phosphorylated STAT3 level was significantly decreased in *IL6ST*^-/-^ THP-1 cells after stimulation with LPS (Figure 5E). The expression of IL-6 was also reduced in *IL6ST*^-/-^ THP-1 cells in response to different inflammatory stimuli (Figure 5F).

**Figure 5.**
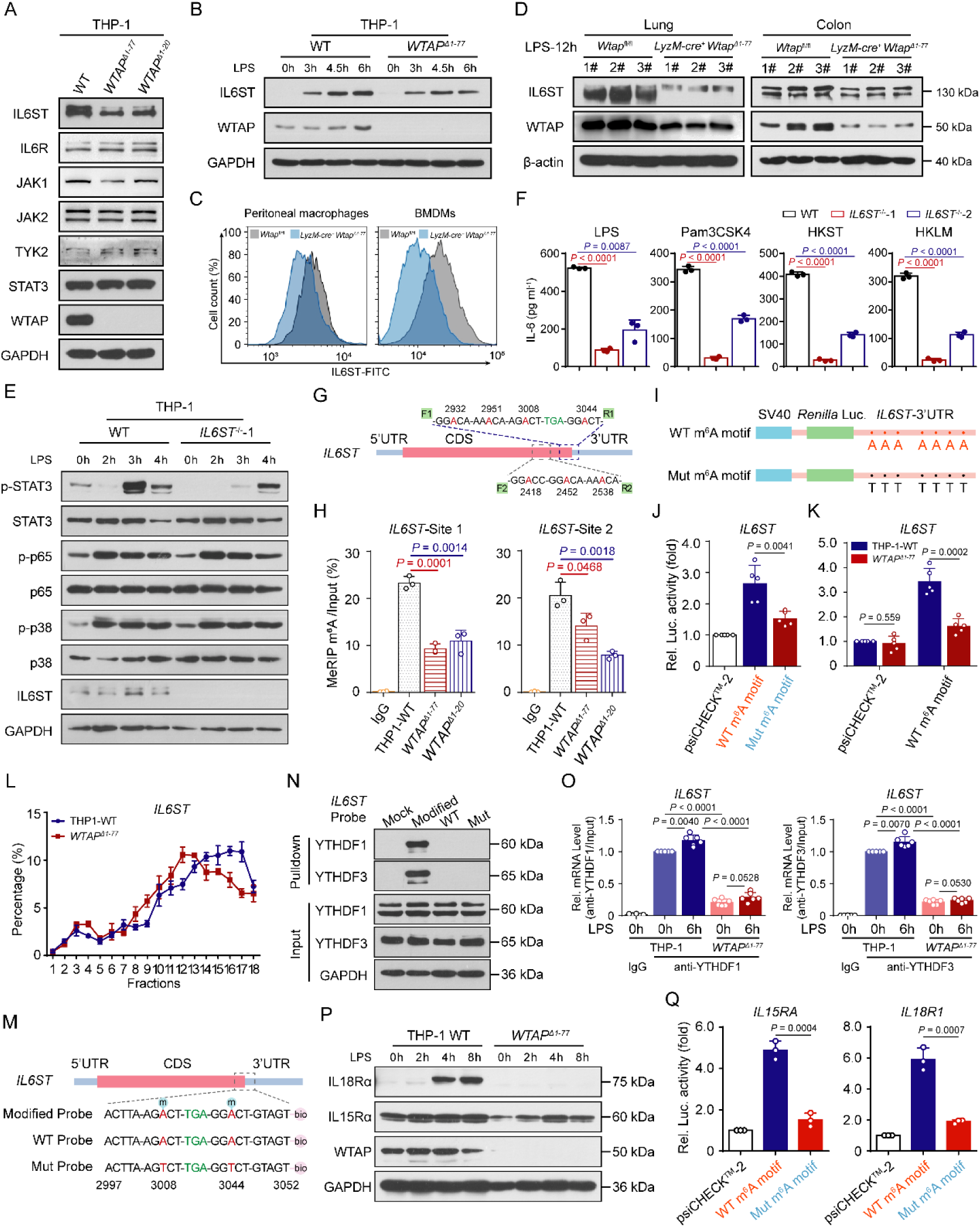
WTAP promotes the protein output of proinflammatory genes through m^6^A modification. (**A**) Immunoblots showing the expression of critical adapters in the IL-6/ STAT3 signaling pathway in WT and *WTAP^Δ1-77^* THP-1 cells. (**B**) Immunoblots showing the expression of IL6ST in WT and *WTAP^Δ1-77^* THP-1 cells stimulated with LPS at the indicated time points. (**C**) Representative flow cytometry data showing IL6ST-FITC fluorescence in cell surface of peritoneal macrophages and BMDMs. (**D**) Immunoblots showing the expression of WTAP and IL6ST in the lung or colon tissues from *Wtap*^fl/fl^ and *LyzM-cre*^+^ *Wtap^Δ1-77^* mice that had been intraperitoneally injected with LPS (10 mg kg^-1^) for 12 hr. (**E**) Immunoblots showing the total and phosphorylated STAT3, p65 and p38 levels in WT and *IL6ST*^-/-^ THP-1 cells stimulated with LPS at the indicated time points. (**F**) ELISAs were performed to measure IL-6 secretion in the supernatants of WT and *IL6ST*^-/-^ THP-1 cells stimulated with LPS, Pam3CSK4, HKST or HKLM for 6 hr. (**G**) Schematic representation showing the position of m^6^A motifs within the IL6ST transcripts. F1/R1 represents detection site 1, which includes four “DRACH” elements, and F2/R2 represents detection site 2, which includes three “DRACH” elements. (**H**) The abundance of *IL6ST* transcripts (with detection sites 1 and 2) among mRNAs immunoprecipitated with an anti-m^6^A antibody in WT and *WTAP^Δ1-77^* THP-1 cells stimulated with LPS for 6 hr. (**I**) WT or mutant (Mut) m^6^A consensus sequences (A-to-T mutation) on the *IL6ST*-3’ UTR were fused with the *Renilla* luciferase reporter in the psiCHECK^TM^-2 vector. (**J**) Relative luciferase activities in 293T cells after transfection with reporter vectors bearing *IL6ST*-3’ UTR with WT or Mut m^6^A sites. *Renilla* luciferase activity was measured and normalized to firefly luciferase activity. (**K**) Relative luciferase activities in WT and *WTAP* KO 293T cells after transfection with reporter vectors bearing the *IL6ST*-3’ UTR with WT m^6^A sites. *Renilla* luciferase activity was measured and normalized to firefly luciferase activity. (**L**) qRT–PCR showing the proportion of *IL6ST* mRNA in polysome fractions from WT and *WTAP^Δ1-77^* THP-1 cells stimulated with LPS for 6 hr. (**M**) Schematic representation of the biotin-labeled probes of *IL6ST* transcripts. (**N**) RNA pull-down analyses showing the interaction between different *IL6ST* RNA probes and the YTHDF1 or YTHDF3 protein. (**O**) YTHDF1 or YTHDF3 were immunoprecipitated, and RNA immunoprecipitation (RIP)-qPCR was performed to assess the association of *IL6ST* with YTHDF1 or YTHDF3 in WT and *WTAP^Δ1-77^* THP-1 cells. (**P**) Immunoblots analyses showing the protein abundance of IL18Rα or IL15Rα in WT and *WTAP*^Δ^*^1-77^* THP-1 cells stimulated with LPS at the indicated time points. (**Q**) Relative luciferase activities in 293T cells after transfection with reporter vectors bearing *IL15RA*- or *IL18R1*-3’ UTR with WT or Mut m^6^A sites. *Renilla* luciferase activity was measured and normalized to firefly luciferase activity. Data are representative of three independent biological experiments in (A to E), (N) and (P). Data are presented as the mean ± s.d. in (F), (H), (J to L), (O) and (Q), with individual measurements overlaid as dots, statistical analysis was performed using a two-tailed Student’s *t*-test.

MeRIP-Seq revealed that the m^6^A peaks obtained with high confidence were distributed in regions near the stop codon in *IL6ST* transcripts (Supplementary Figure 7A). We then designed two gene-specific primer pairs, F1/R1 and F2/R2 (Supplementary Table 3), to measure the change in m^6^A abundance at sites 1 and 2 of *IL6ST* transcripts via MeRIP-qPCR assay (Figure 5G). The results showed that the m^6^A marks enriched at both site 1 and site 2 in the *IL6ST* transcripts were markedly decreased in *WTAP^Δ1-77^* THP-1 cells compared with control cells (Figure 5H). For verification, we mutated the putative m^6^A-modified adenosine by replacing it with a thymine in the *IL6ST* mRNA and inserted the wild-type (WT) or mutated (Mut) UTRs into a reporter gene plasmid (psiCHECKTM-2) (Figure 5I). Luciferase reporter assay data showed that the luciferase activities of the Mut reporter were significantly weaker than those of the WT reporter (Figure 5J). Moreover, the luciferase activities of the WT reporter were weakened in *WTAP^Δ1-77^*THP-1 cells (Figure 5K). Ectopic expression of Flag-tagged WTAP in *WTAP^Δ1-20^* 293T cells enhanced the luciferase activities of the WT reporter (Supplementary Figure 8D). All these data indicated that the detected m^6^A mark sites in *IL6ST* transcripts are direct substrates of WTAP and crucial for maintaining the output of the IL6ST protein.

Because m^6^A modification can affect many aspects of gene expression, including nuclear export, splicing, 3’-end processing, decay and translation (43), we evaluated the mRNA expression of *IL6ST* and found that deleting WTAP did not affect the *IL6ST* mRNA abundance or stability (Supplementary Figure 8, E and F). Due to the decay of *IL6ST* mRNA may not be affected by WTAP, we next measured the translation efficiency of IL6ST through polysome profiling (Supplementary Figure 8G). We calculated the proportion of mRNAs in polysome fractions via qRT‒PCR and found that the distribution of *IL6ST* mRNA shifted to the lighter fraction in *WTAP^Δ1-77^* THP-1 cells after stimulation with LPS for 4 h (Figure 5L). However, this difference was not observed in *GAPDH* mRNA (Supplementary Figure 8H). Taken together, WTAP can facilitate the expression of IL6ST by enhancing the translation efficiency via m^6^A modification. Functional interpretation of m^6^A modification is realized by RNA-binding proteins called “readers”, mainly YTHDF1/2/3 and YTHDC1/2, which can influence the degradation or translation of m^6^A-modified RNA (44, 45). Because YTHDF1 and YTHDF3 are the major m^6^A readers that promote the translation of m^6^A-modified mRNA, we sought to determine whether YTHDF1 or YTHDF3 targets cellular *IL6ST* transcripts to regulate their translation. Immunoblot analyses showed that the deletion of YTHDF1 or YTHDF3 in THP-1 cells using a CRISPR-mediated genome editing approach reduced the protein expression of IL6ST (Supplementary Figure 8I). To verify the direct binding of YTHDF1 and YTHDF3 to m^6^A-methylated *IL6ST* mRNA, we synthesized biotin-labelled RNA probes based on the distribution of the m^6^A peak on *IL6ST* transcripts (Figure 5M) and performed an RNA pull-down assay followed by immunoblotting of the isolated proteins. The results revealed that methylated *IL6ST* transcripts strongly interacted with YTHDF1 and YTHDF3 (Figure 5N). Furthermore, RNA immunoprecipitation (RIP) analyses with antibodies against YTHDF1 or YTHDF3 followed by qRT‒PCR revealed that the amount of *IL6ST* mRNA bound to YTHDF1 or YTHDF3 was significantly increased after stimulation with LPS but reduced after WTAP depletion (Figure 5O). These results suggested that the m^6^A marks on *IL6ST* mRNA mediated by WTAP obviously promoted the binding of YTHDF1 and YTHDF3 to *IL6ST* mRNA. Polysome profiling assays showed that deletion of YTHDF1 or YTHDF3 in THP-1 cells efficiently decreased the protein expression of IL6ST by hindering its translation (Supplementary Figure 8, J and K). Overall, these results demonstrated that the increased efficiency of IL6ST translation induced by the m^6^A modification is mediated by YTHDF1 and YTHDF3.

In addition to IL-6/IL6ST/STAT3 signalling, the IL-15/IL15R and IL18/IL18R signalling which dictate T cell response, regulate B cell homing, and activate NK cells (46), may also mediate the regulation of inflammatory responses by activating STAT3 (47, 48). In our data, the m^6^A peaks distributed in the 3’-UTR regions of *IL15RA* and *IL18R1*, the crucial receptor molecules for IL- 15/IL15R and IL18/IL18R signaling, also almost disappeared when WTAP was depleted (Supplementary Figure 7, B and C). The protein abundance of IL15Rα and IL18Rα was significantly reduced in *WTAP^Δ1-77^* THP-1 cells compared with the control cells (Figure 5P), suggesting that WTAP-mediated m^6^A modification can regulate the expression of IL15Rα and IL18Rα proteins. Further reporter assays performed by replacing the putative m^6^A-modified adenosine with a thymine in the 3’ UTR of *IL15RA* and *IL18R1* mRNA confirmed that the protein abundances of *IL15RA* and *IL18R1* were promoted by WTAP-mediated m^6^A (Figure 5Q and Supplementary Figure 8, L and M). We further rescued the expression of WTAP in *WTAP^Δ1-77^* THP-1 cells and found that the overall m^6^A modification level increased with the ectopic expression of WTAP (Supplementary Figure 8N). Accordingly, the protein abundance of IL6ST, IL15Rα and IL18Rα also increased with elevated m^6^A abundance (Supplementary Figure 8, O and P). All these data revealed that the significance of WTAP for finetuning the activation of IL6/STAT3 signalling axis, implying that the protein synthesis of many proinflammatory genes can be regulated by WTAP via m^6^A modification in immune homeostasis and disease occurrence, which is unique to WTAP compared to other m^6^A regulators.

### The phase separation of WTAP promotes METTL3 recruitment to efficiently modify inflammatory transcripts

By verifying the MeRIP-seq data, we found that the genes affected by WTAP in THP-1 cells did not completely overlap before and after LPS treatment (Supplementary Figure 9A). Transcripts with low m^6^A abundance in *WTAP^Δ1-77^* THP-1 cells were involved in very few metabolic pathways in the resting state (Supplementary Figure 9B) but were mainly enriched in cytokine production and inflammatory signalling pathways after LPS treatment (Supplementary Figure 9C). Additionally, metagene profiles of the m^6^A peak distribution showed that inflammatory stimulation increased the abundance of m^6^A marks but decreased the number of m^6^A peaks (Supplementary Figure 6, C and D). These observations implied that the upregulation of WTAP activated by p65 may lead to more concentrated deposition of m^6^A marks on inflammatory genes.

Recently, liquid-liquid phase separation (LLPS) has emerged as a widespread mechanism through which cells dynamically recruit and organize key signalling molecules. LLPS has also been revealed to play an important role in the formation of the m^6^A “writer” complex and the regulation of the fate of m^6^A-modified mRNAs (49–52). An analysis of the disordered regions in the WTAP protein using the IUPred2A (https://iupred2a.elte.hu/) and PLAAC (http://plaac.wi.mit.edu/) tools revealed a low-complexity region of 16.5 kDa in the C-terminus of the WTAP coding sequence (CDS) (Figure 6, A and B), which may form liquid droplets as a result of phase separation (53). We then further explored whether WTAP has an ability to undergo LLPS upon inflammatory stress and how it plays its roles in the progress of m^6^A modification of those inflammatory transcripts. To this end, recombinantly expressed green fluorescent protein (GFP)-WTAP fusion proteins were bacterially expressed and purified (Supplementary Figure 10A). The GFP-WTAP formed spherical droplets with an aspect ratio close to 1, and an increase in the protein concentration and salt concentration increased the abundance of GFP droplets from barely detectable small foci to regular and large droplets (Figure 6C and Supplementary Figure 10B). Through time-lapse microscopy, we found that the droplet size of GFP-WTAP increased over time (Supplementary Figure 10C), and captured the fusion between droplets within 40 s (Supplementary Figure 10D), implying the high dynamics and fluidity of the GFP-WTAP droplets. Conversely, droplet formation was substantially inhibited by 10% 1,6-hexanediol (Hex), a compound that putatively dissolves liquid-liquid phase-separated condensates (Supplementary Figure 10E).

**Figure 6.**
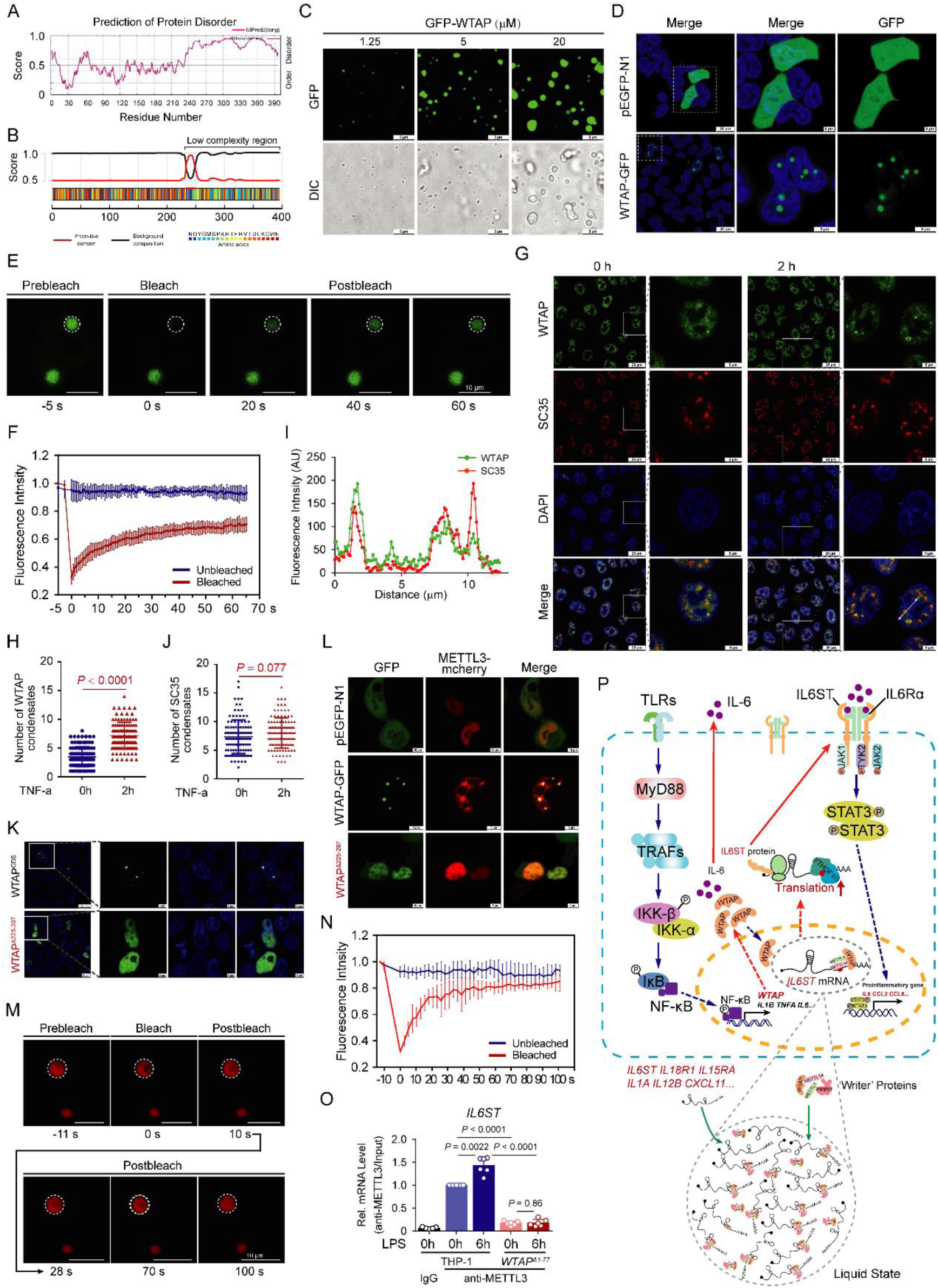
The phase separation of WTAP leads to METTL3 recruitment to modify inflammatory transcripts efficiently. (**A** and **B**) Prediction of the disordered regions and prion-like domains (PrD) of WTAP using the IUPred2A (A) or PLAAC (B) tool. (**C**) Images of GFP- WTAP droplets formation at room temperature with indicated GFP-WTAP concentrations (350 mM NaCl). (**D**) LSCM images of 293T cells expressing PEGFP-N1 or GFP-tagged WTAP constructs. (**E**) FRAP assays of the GFP-tagged WTAP droplets before and after photobleaching. (**F**) FRAP quantification of GFP-WTAP droplets over a period of 65 s. (**G**) LSCM images of HeLa cells treated with TNF-α for 2 hr. (**H**) Statistics of the number of WTAP droplets in (G). **(I)** Quantitative line profile of colocalization along a white arrow of the image of (G). (**J**) Statistics of the number of SC35 droplets in (G). (**K**) LSCM images of 293T cells expressing full-length or truncated WTAP labeled with GFP. (**L**) LSCM images of the mCherry-tagged METTL3 droplets (red) in 293T cells before and after co-transfection with GFP-tagged full-length or truncated WTAP. (**M**) FRAP assays of the mCherry-tagged METTL3 droplets before and after photobleaching from (L). (**N**) FRAP quantification of mCherry-METTL3 droplets over a period of 100 s. (**O**) METTL3 was immunoprecipitated and RIP-qPCR was performed to assess the association of *IL6ST* transcripts with METTL3. (**P**) Working model for WTAP facilitating inflammatory responses through m^6^A modification and phase separation. Data are representative of three independent biological experiments in (C to E), (G), and (K to M). Data are presented as the mean ± s.d. in (H), (J) and (O), with individual measurements overlaid as dots, statistical analysis was performed using a two-tailed Student’s *t*-test. Indicated scale bars are shown in (C to E), (G), and (K to M).

To further test the ability of WTAP to undergo LLPS in intact cells, we constructed a C-terminal enhanced GFP (EGFP)-tagged WTAP vector and ectopically expressed it in 293T cells (Supplementary Figure 10F). We found that GFP-tagged WTAP formed discrete puncta in the nucleus (Figure 6D). In line with those in vitro results, fluorescence recovery after photobleaching (FRAP) assays in 293T cells showed that the fluorescence of GFP-tagged WTAP was efficiently and gradually recovered after bleaching (Figure 6, E and F), indicating the potential phase separation capability of WTAP in vivo. Notably, endogenous WTAP condensates formed in HeLa cells, and the number of these droplets increased significantly after cell stimulation with TNF-α due to upregulated expression of WTAP (Figure 6, G and H and Supplementary Figure 10G), suggesting that upregulated WTAP induced by inflammatory stimuli is more prone to phase separation. In consistent with other study, the WTAP droplets colocalized with the nuclear speckle marker SC35 (19) (Figure 6, G and I). However, no difference in the number of SC35 droplets was found before and after inflammatory stimulation (Figure 6J), suggesting that endogenous WTAP droplets formed by self-occurring LLPS rather than the dynamics of the nuclear speckles, and the LLPS of WTAP may facilitate its colocalization with nuclear speckles.

To further identify the regions in WTAP that are needed for phase separation, we used the PONDR tool (http://www.pondr.com) to predict 5 unfolded intrinsically disordered regions (IDRs) with scores greater than 0.7 in the WTAP protein sequence (Supplementary Figure 10H). A series of truncated WTAP mutants labelled with GFP were then constructed (Supplementary Figure 10H) and expressed in 293T cells (Supplementary Figure 10I). Through confocal imaging, we found that WTAP failed to form droplets only when amino acid residues 225-287 were deleted (Figure 6K and Supplementary Figure 10J), suggesting that this sequence is essential for the LLPS of WTAP. Biomolecular condensates formed through LLPS can maintain locally elevated concentrations of resident proteins and/or RNAs (54). Hence, we subsequently explored the biological function of the phase separation of WTAP. Previous studies revealed that WTAP recruits METTL3 and other cofactors to nuclear speckles and that the RNA-binding capacity of METTL3 is profoundly reduced in the absence of WTAP (19). Here, we found that mCherry-METTL3 (Supplementary Figure 10K) readily fused with phase-separated GFP-WTAP and colocalized to all GFP-WTAP droplets (Supplementary Figure 10L), demonstrating that fusion with WTAP enhanced the METTL3 LLPS in vitro. In vivo, METTL3 also formed liquid droplets with the help of WTAP (Figure 6, L-N), since deleting the LLPS sequence on WTAP significantly reduced the ability of WTAP to recruit METTL3 through LLPS (Figure 6L) without affecting the direct binding of METTL3 and WTAP (Supplementary Figure 10M), suggesting that WTAP LLPS might prime METTL3 condensation in cells. To confirm that the LLPS of WTAP promotes METTL3 to methylate inflammatory mRNAs, RIP with antibodies against METTL3 followed by qRT‒PCR was performed and the results revealed that the amount of *IL6ST*, *IL15RA* and *IL18R1* mRNAs bound by METTL3 was significantly increased after stimulation with LPS and was reduced by WTAP depletion (Figure 6O and Supplementary Figure 10, N and O). These data further indicated that the LLPS of WTAP facilitates the aggregation of METTL3 and the methylation of inflammatory mRNAs. Consistently, the absence of phase transition capability observably inhibited WTAP-dependent m^6^A modification (Supplementary Figure 10P). Taken together, these data suggested that upregulated WTAP undergoes phase separation to facilitate the assembly of the writer complex and the localization of the writer complex to nuclear speckles. Since the nuclear speckles are active regions for gene transcription and RNA processing and are rich in transcriptionally active proinflammatory transcripts during inflammatory stress, WTAP-medicated LLPS may improve the accessibility between the m^6^A writer complex and inflammatory transcripts in nuclear speckles, promoting the deposition of m^6^A onto transcriptionally active inflammatory transcripts and the activation of proinflammatory responses (Figure 6P).

### WTAP deficiency alleviates the disease progression of DSS-induced IBD in mice

All the aforementioned data suggest that WTAP is a true proinflammatory risk factor. To further investigate the potential role and clinical relevance of upregulated WTAP in the progression of inflammatory diseases, we first revealed that both WTAP and IL6ST were upregulated in SLE, asthma, sepsis, RA, psoriasis and IBD patients (Supplementary Figure 1, A-E and Supplementary Figure 11A). Moreover, high expression of WTAP was significantly positively correlated with the level of inflammation in patients. For example, the upregulation of WTAP was significantly inhibited in psoriasis patients treated with anti-IL17A therapy and Crohn’s disease (CD) patients treated with anti-TNF therapy (GSE137218/16879; Supplementary Table 1), correspondingly, proinflammatory cytokine expression was also significantly decreased in these patients after treatment (Figure 7, A and B). We further evaluated the expression of WTAP in the progression of IBD. An analysis of a single-cell dataset of IBD patients (GSE125527; Figure 7C and Supplementary Table 1) showed that the expression of WTAP rather than METTL3 was significantly increased in some immune cell subsets in IBD patients, particularly in monocytes (Figure 7D). The prevalence of WTAP upregulation in IBD patient samples was then verified using more public datasets (GSE119600/10616; Supplementary Figure 11, B and C and Supplementary Table 1). Moreover, using data from the IBDMDB database, an analysis of the correlation between genes involved in m^6^A methylation and genes that have been identified as IBD risk genes or biomarkers showed that WTAP exhibited the highest correlation with the expression of IBD genes among all the m^6^A regulators (Supplementary Figure 11D). Observably, increased expression of WTAP was accompanied by an increase in the global m^6^A abundance in various inflammatory diseases (Supplementary Figure 11, E and F), which is consistent with our results in LPS-induced macrophages. Consistent with this finding, similar tendencies were observed in mouse models of imiquimod (IMQ)-induced psoriasis (Figure 7E) and 3% dextran sodium sulfate (DSS)-induced colitis (Figure 7F). Thus, the high expression of WTAP accompanied by a high level of m^6^A modification and high expression levels of proinflammatory cytokines should be a common characteristic of distinct inflammatory diseases, such as IBD.

**Figure 7.**
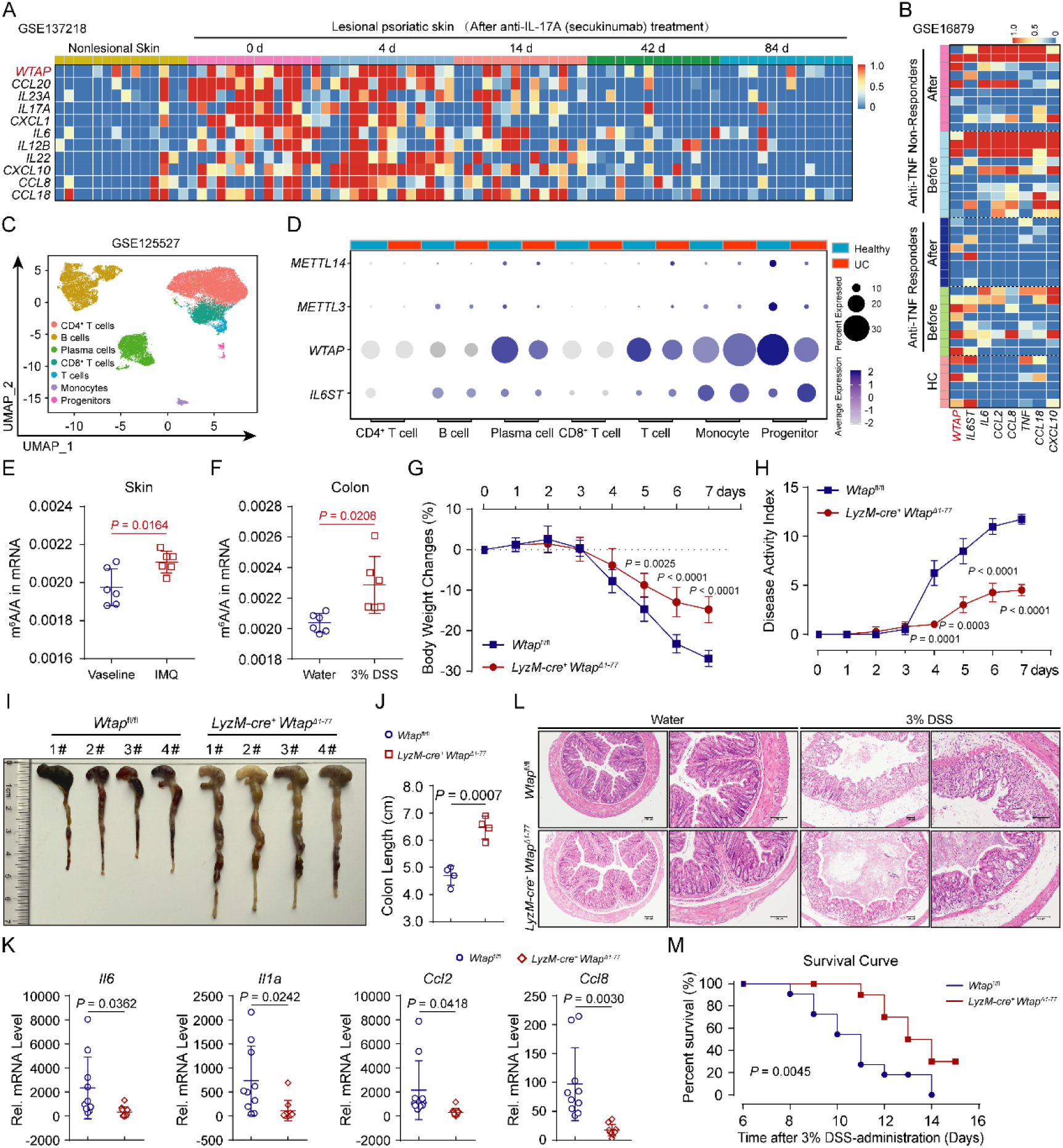
WTAP deficiency alleviates the disease progression of DSS-induced IBD in mice. (**A**) Heatmap showing the expression of WTAP and proinflammatory genes in samples from psoriasis patients before and after anti-IL-17A therapy. (**B**) Heatmap showing the expression of WTAP and proinflammatory genes in samples from CD patients before and after anti-TNF therapy. (**C**) Uniform manifold approximation and projection (UMAP) plots showing rectal tissue-derived CD45^+^ immune cells from all participants. The meaning of the cluster colors is indicated. (**D**) Dot plots showing the expression of METTL14, METTL3, WTAP and IL6ST in major immune cell groups. (**E** and **F**) LC-MS/MS quantification of m^6^A abundance in mRNA extracted from skin tissues in IMQ-induced psoriasis (**E**) or colon tissues in DSS-induced colitis (**F**) of mice. (**G**) Body weight changes of mice given 3% DSS in their drinking water were monitored daily. n = 7 mice per group. (**H**) The disease activity index of *Wtap*^fl/fl^ and *LyzM-cre*^+^ *Wtap^Δ1-77^* mice was scored daily. n = 7 mice per group. (**I** and **J**) Macroscopic appearances (I) and colon lengths (J) of *Wtap*^fl/fl^ and *LyzM- cre*^+^ *Wtap^Δ1-77^* mice were recorded on day 6. n = 4 mice per group. (**K**) qRT‒PCR showing the mRNA abundance of *Il6*, *Il1α*, *Ccl2* and *Ccl8* in the colon tissues from *Wtap*^fl/fl^ and *LyzM-cre*^+^ *Wtap^Δ1-77^* mice that had been given 3% DSS in their drinking water for 6 days. n = 10 mice per group. (**L**) Histopathological changes in colon tissue were determined by H&E staining. Scale bars, 100 μm. n = 4 mice per group. (**M**) The survival of the *Wtap*^fl/fl^ and *LyzM-cre*^+^ *Wtap^Δ1-77^* mice after continuous feeding with 3% DSS was monitored for 15 days. n = 10 mice per group. Data are representative of three independent biological experiments in (I) and (L). Data are presented as the mean ± s.d. in (E), (F), (G and K), with individual measurements overlaid as dots, statistical analysis was performed using a two-tailed Student’s *t*-test. Indicated scale bars are shown in (I) and (L).

To verify the biological correlation between WTAP and IBD signatures, we subjected *LyzM*-Cre^+^ *Wtap^Δ1-77^* mice and their *Wtap*^fl/fl^ control littermates to DSS to induce acute colitis. The administration of DSS in drinking water can cause the death of intestinal epithelial cells and thus compromise gut barrier function and cause inflammation (55). We found that compared with their *Wtap*^fl/fl^ control littermates, the *LyzM*-Cre^+^ *Wtap^Δ1-77^*mice treated with DSS displayed attenuated colitis, accompanied by less weight loss (Figure 7G) and a lower disease activity index (Figure 7H). Moreover, WTAP deficiency substantially attenuated the shortening of the colon length in the DSS- challenged mice (Figure 7, I and J). The low expression of *Il6*, *Il1α*, *Ccl2* and *Ccl8* in DSS- challenged *LyzM*-Cre^+^ *Wtap^Δ1-77^*mice indicated attenuated colonic inflammation (Figure 7K). A histopathological assessment revealed that the colonic mucosa of *LyzM*-Cre^+^ *Wtap^Δ1-77^* mice was relatively intact, with less inflammatory cell infiltration after DSS treatment (Figure 7L).

Furthermore, *LyzM*-Cre^+^ *Wtap^Δ1-77^* mice exhibited later death onset and a lower death rate (Figure 7M). Taken together, these results suggested that WTAP deficiency attenuates the severity of DSS-induced IBD, revealing a clinically relevant correlation between the WTAP level and IBD progression. Because the increased expression of WTAP is ubiquitous in many inflammatory diseases, we hypothesize that WTAP is a novel and important risk factor that exerts a broad spectrum of effects on inflammatory diseases and may serve as a novel potential therapeutic target in the treatment of inflammatory diseases.

### Reducing the level of m^6^A modification can reverse the high inflammatory state

As we have shown above, hyperinflammatory states are often accompanied by high levels of m^6^A mark, we further test whether STM2457, a METTL3 inhibitor, can alleviate hyperinflammatory states by reducing m^6^A modification. The results showed that reducing the abundance of m^6^A marks by STM2457 (Figure 8A) significantly inhibited HKST-induced expression of proinflammatory genes (Figure 8B). We also verified this effect in a mouse model of LPS-induced sepsis by showing that the m^6^A modification levels of colon and lung tissues from mice were significantly reduced by STM2457 (Figure 8, C and D). And a decrease in the m^6^A modification level blocked the up- regulation of inflammatory cytokines and activation of inflammatory signals induced by LPS (Figure 8, E and F). Similarly, STM2457 effectively alleviated lung inflammation and the pathological characteristics of lung injury (Figure 8G). Thus, reducing the global m^6^A levels may be a potential therapeutic strategy for alleviating hyperinflammation.

**Figure 8.**
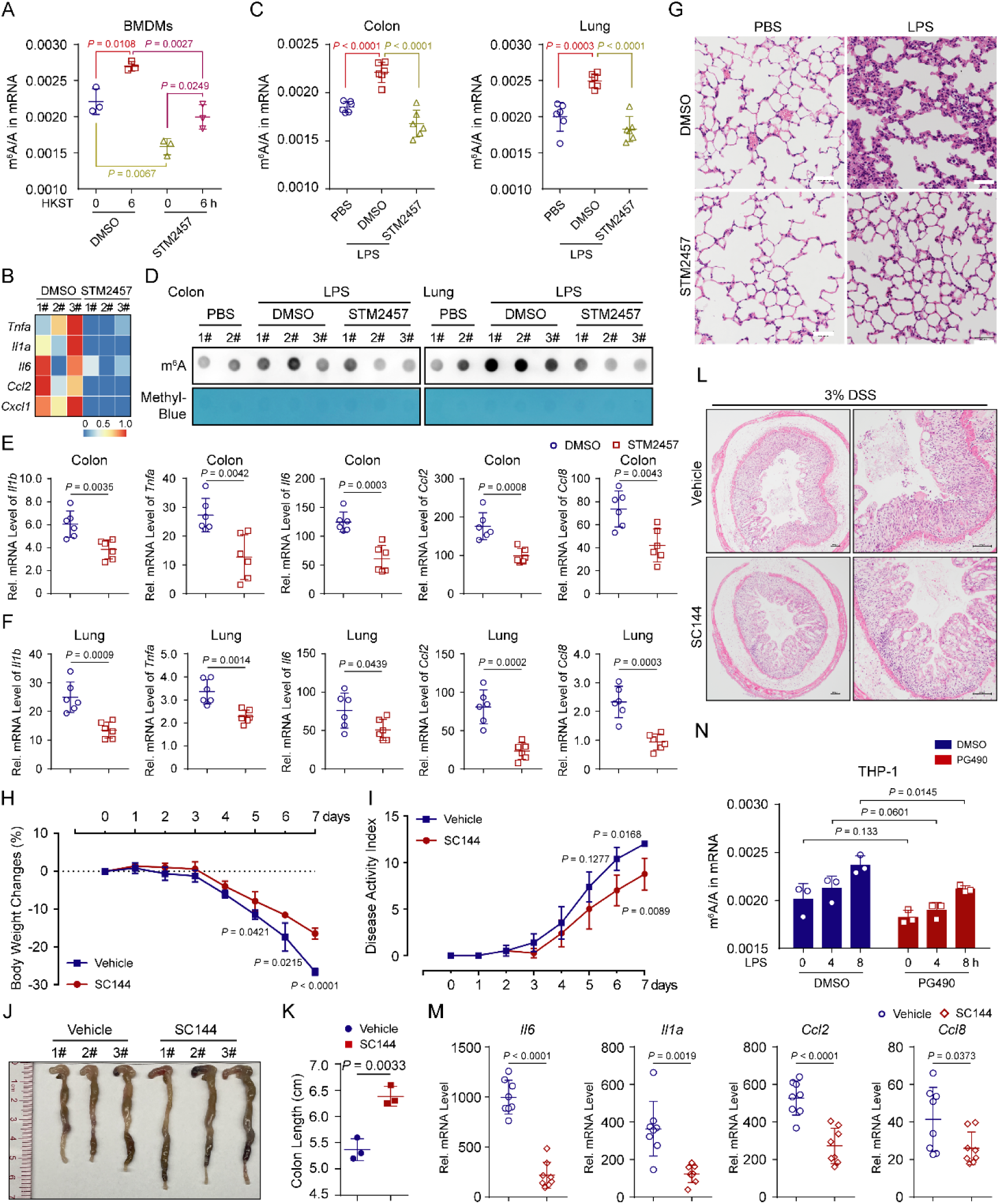
Reducing the level of m^6^A modification can reverse the high inflammatory state. (**A**) LC–MS/MS quantification of m^6^A abundance in mRNA extracted from BMDMs that pretreated with DMSO or STM2457, followed by stimulation with HKST for 6 hr. UN, unstimulated. (**B**) Heatmap showing the mRNA expression of proinflammatory genes in BMDMs pretreated with DMSO or STM2457, followed by stimulation with HKST for 6 hr. The number represents three independent experiments. (**C** and **D**) LC–MS/MS (C) and m^6^A dot blot (D) quantification of m^6^A levels in mRNA extracted from colon or lung tissues in LPS-induced sepsis of mice, followed by four consecutive days of intraperitoneal injections of DMSO or STM2457. n = 3 mice per group. (**E** and **F**) qRT‒PCR showing the mRNA abundance of *Il1b*, *Tnfa*, *Il6*, *Ccl2* and *Ccl8* in the colon (E) or lung (F) tissues from mice treated as above. n = 6 mice per group. (**G**) H&E assays showing the lung injury of LPS-induced sepsis of mice, followed by four consecutive days of intraperitoneal injections of DMSO or STM2457. Scale bars, 50 μm. n = 3 mice per group. (**H**) Body weight changes of mice intraperitoneally injected with vehicle or SC144, followed by drinking with 3% DSS water were monitored daily. n = 4 mice per group. (**I**) The disease activity index of mice intraperitoneally injected with vehicle or SC144, followed by drinking with 3% DSS water was scored daily. n = 4 mice per group. (**J** and **K**) Macroscopic appearances (J) and colon lengths (K) of mice were recorded on day 8. n = 3 mice per group. (**L**) Histopathological changes in colon tissue were determined by H&E staining. Scale bars, 100 μm. n = 3 mice per group. (**M**) qRT‒PCR showing the mRNA abundance of *Il6*, *Il1α*, *Ccl2* and *Ccl8* in the colon tissues from mice intraperitoneally injected with vehicle or SC144, and then given 3% DSS in their drinking water for 8 days. n = 8 mice per group. (**N**) LC–MS/MS quantification of m^6^A levels in mRNA extracted from THP-1 cells that treated with DMSO or PG490, followed by stimulation with LPS at the indicated time points. Data are representative of three independent biological experiments in (D), (G) and (J). Data are presented as the mean **± s.**d. in (A), (C), (E and F), (K) and (M and N), with individual measurements overlaid as dots, statistical analysis was performed using a two-tailed Student’s *t*-test.

In addition, the specific IL6ST inhibitor Bazedoxifene (BZA) has long been approved by the US Food and Drug Administration as a selective modulator of oestrogen receptor to treat osteoporosis (56), hinting at the promise of IL6ST inhibitors as a drug. In this study, we found that WTAP positively regulates proinflammatory response primarily through IL6ST. So, is blocking IL6ST a promising attempt to treat inflammatory bowel disease or even other inflammatory diseases in the future? To test this hypothesis, we injected the vehicle or SC144 intraperitoneally, along with feeding 3% DSS drinking water, and monitored the disease severity of mice daily. The results showed that compared with control littermates, the mice treated with SC144 displayed attenuated colitis, accompanied by less weight loss (Figure 8H) and a lower disease activity index (Figure 8I).

Moreover, SC144 treatment attenuated the shortening of the colon length in the DSS-challenged mice (Figure 8J and K). A histopathological assessment (Figure 8L) and the low expression of *Il6*, *Il1α*, *Ccl2* and *Ccl8* in SC144-treated mice indicated attenuated colonic inflammation (Figure 8M). These data suggested that blocking the activation of STAT3 signals by inhibiting IL6ST can alleviate the disease progression of DSS-induced IBD in mice. In line with this, a recent study also pointed out that gp130 (IL6ST) blockade may benefit some patients with Crohn’s disease (57). Hence, our study confirmed that screening suitable inhibitors with few side effects to block IL6ST maybe a promising attempt to treat inflammatory bowel diseases in the future.

## Discussion

### WTAP acts as a smart adapter in the assembly of the m^6^A writer complex to participate in inflammation regulation

Inflammation is vital for protecting the host against invading pathogens and for repairing tissue damage, which requires tight and concise control of pro- and anti-inflammatory gene expression. m^6^A, the most prevalent internal mRNA modification, has been recently linked to various inflammatory states, including autoimmunity, infection, metabolic diseases and cancers. In this study, we further found that WTAP, as a p65-controlled gene, is universally upregulated under distinct inflammatory stimuli. Increased WTAP protein is strongly related to higher levels of m^6^A modification and excessive inflammatory responses. Noteworthy, the transcription and expression of WTAP have been shown to be regulated in various biological processes. For example, at the transcriptional levels, HIF1α (37) and STAT3 (36) can transactivate WTAP gene expression by directly binding to its promoter region, and WTAP can also be upregulated by the epigenetic alteration of H3K4me3 (58). At the translational levels, pseudogene WTAPP1 can bind to WTAP mRNA to promote its translation by recruiting the EIF3 translation initiation complex, accelerating the progression of pancreatic ductal adenocarcinoma (PDAC) (59). At the posttranslational levels, our previous study showed that WTAP undergoes degradation via the ubiquitination-proteasome pathway in virus-infected cells, leading to reduction in the abundance of m^6^A marks and the subsequent attenuation in the intensity of IFN-I signalling (60). Here, we further found that WTAP can also be significantly activated by NF-κB p65 and slightly activated by C/EBPβ under inflammatory stress. And unlike other m^6^A proteins that regulate inflammatory responses highly dependent on context, WTAP is more commonly involved in the establishment and occurrence of hyperinflammatory states and inflammatory diseases. In short, using smart adapters without methyltransferase activity, such as WTAP, may provide more precise adjustment of m^6^A “writer” complex activity, which is evolutionarily and physiologically beneficial to the dynamic regulation of phenotypic diversity and homeostasis in higher organisms.

In addition to showing that WTAP is a p65-controlled gene, the basic mechanism by which WTAP coordinates the assembly dynamics of the writer complex to promote the m^6^A modification of many inflammatory genes was further revealed in this study. Although WTAP has been shown to be critical for the localization of the m^6^A “writer” complex to nuclear speckles, its roles in the dynamic assembly of the “writer” complex is not fully unexplored. A previous study reported that METTL3 can undergo LLPS (52), and METTL3 interacts with m^6^A-METTL-associated complex (MACOM) mainly through WTAP (61). Here, we further revealed that highly expressed WTAP spontaneously forms condensates through LLPS under inflammatory stress. Moreover, when WTAP is absent or cannot undergo LLPS, METTL3 is dispersed in the nucleus, indicating that the LLPS of WTAP is essential for the assembly of the writer complex and provides dynamic m^6^A regulation under distinct physiological or pathological conditions, which may be important for the recruitment of more “writer” proteins to nuclear speckles. Because nuclear speckles are sites with many transcriptionally active genes (62), WTAP-mediated LLPS may increase the local “writer” concentrations and improve the accessibility between the m^6^A “writer” complex and inflammatory transcripts in nuclear speckles, thereby promoting the m^6^A modification of inflammatory genes and accelerating of inflammatory response. In the past decade, biomolecular condensates formed by LLPS have been widely reported to modulate many cellular functions by compartmentalizing specific proteins and nucleic acids in subcellular environments with distinct properties (63, 64). Our finding further expands the regulatory roles of phase separation to the dynamic assembly of writer complex and the methylation of specific transcripts, describing how cells utilize the composition and compartmentation of multivalent condensates to affect the m^6^A epitranscriptome.

### The finetuning of the STAT3 signalling axis by targeting WTAP LLPS may be an effective therapeutic strategy for excessive inflammation

m^6^A modification has been reported to regulate inflammatory responses and related diseases. For example, KDM6B (28), SOCS1 (65), TRAF3 and TRAF6 (24, 66) are dynamically regulated by m^6^A modification in controlling inflammatory responses. Through analyses of clinical data and mouse disease models, we clearly found that increased expression of WTAP is associated with inflammatory diseases, such as sepsis, SLE, asthma (67), RA, psoriasis (68) and IBD (69) (Supplementary Figure 1). A recent study also suggested the relationship of WTAP to the anti-inflammatory effect of *Astragalus mongholicus polysaccharide* (APS) (70). In addition to revealing the relationship between increased WTAP and higher m^6^A modification in inflammation, we specifically showed that an increase in WTAP protein was accompanied by increases in the m^6^A abundances on *IL6ST*, *IL15RA*, *IL18R1* and many other transcriptionally active proinflammatory cytokines. High IL6ST, IL15RA and IL18R1 protein abundance promotes the activation of the STAT3 signalling axis, which in turn contributes to the production of proinflammatory cytokines, such as IL-6, CCL2 and CCL8 (Figure 6P). Consistently, two recent studies individually noted that the knockdown of FTO can significantly increase the overall m^6^A abundance, the phosphorylation of STAT3 and the secretion of proinflammatory cytokines and thereby positively regulates the inflammation or IFN-I response (26, 71). Moreover, the IL-6/STAT3 pathway is aberrantly hyperactivated in many cancers, and such hyperactivation is generally associated with a poor clinical prognosis (72). Recent studies have also shown that high levels of m^6^A modification enhance JAK1 translation in high-metastatic hepatocellular carcinoma (HCC) cells, thereby promoting the activation of STAT3 (73). Interestingly, the expression of WTAP is also significantly elevated in many cancers, such as pancreatic carcinoma (59), osteosarcoma (74), HCC (75) and nasopharyngeal carcinoma (76). This implies that the strategy of finetuning the STAT3 signalling axis by WTAP may also play an important role in the progression of multiple cancers. Because the STAT3 axis has been well characterized as a mechanism to accelerate the production of a variety of cytokines and chemokines (77) and is highly associated with inflammatory diseases, including cytokine storm syndromes, autoimmune diseases and cancers (78, 79), the novel NF- κB/WTAP/STAT3 axis identified in this study indicated that WTAP is an ideal therapeutic target in the treatment of many inflammatory diseases and cancers.

To date, effective therapeutic interventions for excessive inflammation by inhibiting NF-κB or STAT3 activity in the clinic remain to be further developed, largely due to that the challenge of directly targeting the NF-κB and STAT3 causes side effects that affect the homeostasis of the immune system and normal cell growth, differentiation and survival (13, 14). Here, we found that hyperinflammatory states in the body are often accompanied by high levels of m^6^A methylation, and reducing the abundance of m^6^A marks by STM2457 can observably reverse the hyperinflammation. Based on these data combined with one study showing that STM2457 can effectively reduce the severity of DSS-induced IBD in mice (80), we believe that reversing the high m^6^A levels in disease states has broad prospects in the treatment of inflammatory diseases in the future. Because WTAP is more concentrated in the inflammatory context to affect the m^6^A modification of nascent inflammatory genes, the epigenetic therapies by targeting WTAP to reprogram the m^6^A landscape in cells may achieve more selective and safer therapeutic outcomes.

In addition, we found that the NF-κB p65 inhibitor PG490 significantly reduced the overall m^6^A abundance by inhibiting the expression of WTAP (Figure 8N). Thus, for the development of novel small anti-inflammatory molecules, researchers may need to consider the effects of these inhibitors on epitranscriptome.

In summary, we provide the first demonstration that high m^6^A modification in a variety of hyperinflammatory states is p65-dependent, due to that the key component of the writer complex WTAP is transcriptionally regulated by p65 and its overexpression can lead to higher m^6^A modification. We also discovered that upregulated WTAP undergoes phase separation, which facilitates the aggregation of the writer complex and its localization to nuclear speckles and the deposition of m^6^A onto transcriptionally active transcripts, resulting in the promotion of proinflammatory responses and the exacerbation of inflammatory diseases. Hence, we hypothesize that the upregulation of WTAP should be a risk factor in many hyperinflammatory states and inflammatory diseases. Interrupting the assembly of the m^6^A writer complex by targeting the phase separation of WTAP to reduce the global m^6^A level may be a potential therapeutic strategy for preventing excessive inflammation.

## Methods

### Sex as a biological variable

Our study examined male and female animals, and similar findings are reported for both sexes.

### Mice

C57BL/6 *LyzM*-Cre^+^ *Wtap^Δ1-77^* mice were generated by Gempharmatech Co., Ltd. using the CRISPR/Cas9 approach. All mouse lines were maintained at Sun Yat-sen University under specific pathogen-free (SPF) conditions in ventilated microisolator cages. The Institutional Animal Care and Use Committee (IACUC) of Sun Yat-sen University (Guangzhou, China) approved all the experimental protocols concerning the handling of mice. The study is compliant with all relevant ethical regulations regarding animals.

### Cells

The 293T, HeLa and THP-1 cell lines were purchased from ATCC. All these cells were cultured in endotoxin-free DMEM or RPMI-1640, supplemented with 10% FBS and 1% penicillin– streptomycin. PBMCs and BMDMs were isolated and cultured as described previously (60).

### Reagents

Reagents used are listed in Supplementary Table 4.

### In vitro recombinant protein expression and purification

The expressing plasmids (pET-32a) encoding WTAP conjugated eGFP and METTL3 conjugated mCherry were transformed into BL21 *E.coli*. *E.coli* were cultured in LB with Ampicillin (50 µg/mL) at 37 °C for about 12 hr till OD600 = 0.6. After induction with 1 mM IPTG at 37 °C for 8 hr, the cultured *E.coli* was harvested by centrifugation at 4000 rpm, 4 °C, 10 min and resuspended in lysis buffer. Cells were lysed by sonication on ice and centrifuged (12,000 rpm, 30 min, 4 °C) to remove debris and collected the supernatant. The supernatant was purified by incubation with Ni-NTA agarose beads (QIAGEN) overnight at 4 °C. Then, Ni-NTA beads were washed with wash buffer (50 mM NaH2PO4, 300 mM NaCl, 20 mM imidazole, pH 7.8), and proteins were eluted with elution buffer (50 mM NaH2PO4, 300 mM NaCl, 300 mM imidazole, pH 7.8). The purified proteins were further dialyzed by using PD10 column (GE Healthcare), and concentrated using Amicon Ultra 30 K (Millipore) concentrators at 4 °C. Concentrated proteins were quantified by the BCA method (Thermo Fisher) and stored at -80 °C.

### Generation of knockout THP-1 cells

Knockout (KO) cells were constructed using the CRISPR/Cas9 system. Small guide RNAs (gRNAs) targeting the genome sequence of target genes were designed using an online gRNA design tool (by Zhang Feng laboratory, http://crispr.mit.edu/), and subcloned into the lentiCRISPR v2 vector. This vector was transfected into the HEK293T cells along with the following two lentiviral packing plasmids: psPAX2 and pVSV-G. The culture supernatants were collected at 48 and 60 hours after transfection and concentrated by ultracentrifugation before use for infection. The infection-positive (GFP^+^) cells were selected by flow cytometry, monoclonal cells were confirmed by genomic sequencing and immunoblotting analysis with the corresponding antibody. The gRNA sequences used for generating the KO cells are listed in Supplementary Table 3.

### In vivo LPS challenge

For endotoxicity studies, age-matched *Wtap*^fl/fl^ and *LyzM-cre*^+^ *Wtap^Δ1-77^* mice (8 weeks old) were intraperitoneally injected with LPS (40 mg kg^-1^). Mouse survival was monitored every 4 hr.

*Wtap*^fl/fl^ and *LyzM-cre*^+^ *Wtap^Δ1-77^*mice (8 weeks old) were intraperitoneally injected with LPS (10 mg kg^-1^) or isodose PBS. After 12 hr, the mice were killed, blood was collected, and serum levels of IL-6 were measured by ELISA. Lung and colon tissues were collected, and RNA samples were extracted for qRT-PCR to determine the mRNA levels of inflammatory cytokines.

*Wtap*^fl/fl^ and *LyzM-cre*^+^ *Wtap^Δ1-77^*mice (8 weeks old) were intraperitoneally injected with LPS (10 mg kg^-1^) or PBS. After 6 hr, the mice were killed, and lungs from control or LPS-stimulated mice were dissected, fixed in 4% paraformaldehyde (PFA) embedded in paraffin, sectioned, stained with hematoxylin and eosin (H&E) solution, and examined by light microscopy to detect histological changes.

*Wtap*^fl/fl^ and *LyzM-cre*^+^ *Wtap^Δ1-77^*mice (8 weeks old) were intraperitoneally injected with *P. aeruginosa* (ATCC27853, 2 × 10^8^ CFU) or *L. monocytogenes* (ATCC19116, 1 × 10^8^ CFU) or PBS. After 16 hr, the mice were killed, and the related phenotypes were measured as described above.

### ELISA assays and quantitative RT–PCR (qRT–PCR)

The concentrations of IL-6, IL-1β and TNF-α in culture supernatants were measured using kits from R&D Systems or Proteintech, according to the manufacturer’s instructions.

Total RNA was extracted using TRIzol reagent (Invitrogen) and reversed-transcribed with a PrimeScript^TM^ RT reagent kit with gDNA Eraser (TaKaRa) according to the manufacturer’s instructions. And 2×Polarsignal^®^ qPCR mix (MIKX) was used for quantitative real-time PCR analysis. The data were normalized by the level of *GAPDH* or *ACTB* expression in each individual sample, and primer sequences used are listed in Supplementary Table 3.

### Luciferase reporter gene assay

For a promoter reporter gene assay, 293T cells were plated in 48-well plates and transiently transfected with WTAP promoter reporter (pGL3 basic plasmid), with increasing doses of Flag-tagged NF-κB p65, IRF3 or C/EBPβ and 7.5 ng of the *Renilla* luciferase reporter vector using Lipofectamine 2000. At 24 hrs post-transfection, luciferase activities were measured with a dual-luciferase reporter assay system (Promega) according to the manufacturer’s instructions. Reporter gene activity was determined by normalization of Firefly luciferase activity to *Renilla* luciferase activity.

For other reporter gene assays, wild-type and *WTAP^Δ1-20^* 293T cells or wild-type *WTAP^Δ1-77^* THP-1 cells were plated in 48-well plates and transiently transfected with the indicated reporter vectors (psiCHECK^TM^-2) expressing wild-type or mutated UTRs of *IL6ST* using Lipofectamine 2000. At 24 hrs post-transfection, the luciferase activities were measured as above. Reporter gene activity was determined by normalization of the *Renilla* luciferase activity to Firefly luciferase activity.

### RNA decay assay

*WTAP^Δ1-77^* and WT THP-1 cells were seeded at a density of 8 × 10^5^ cells/mL in 12-well plates. After treatment with PMA (50 ng/mL, Sigma), the cells were treated with transcription inhibitor Act D (5 μg/mL, Sigma) to block *de novo* RNA synthesis, and collected at different times. RNA samples were extracted for qRT‒PCR to determine the mRNA levels of the indicated genes.

### Laser scanning confocal microscopy (LSCM)

Cells for fluorescence experiments were cultured in glass bottom culture dishes (Nest Scientific). After treated with indicated stimulation, cells were fixed by 4% paraformaldehyde for 10 min, followed with 3 times wash with PBS. Then, cells were permeabilized with methyl alcohol for 30 min at −20 °C and rinsed with PBS for 3 times. After blocking in 5% goat serum for 1 h at room temperature, cells were incubated with primary antibodies at 4 °C, overnight. 1 × PBST (PBS with 0.1% Tween20) was used to wash the cells for 3 times and subsequently incubated with fluorescently labeled secondary antibodies at room temperature for 1 h. Confocal images were obtained by Leica TCS SP8 STED 3X confocal microscope. The number of condensates were counted and analyzed by ImageJ software.

### Live-cell imaging

At 24 hrs post-transfection incubation, cells were loaded into temperature- and CO2-controlled live-cell imaging chamber of Leica TCS SP8 STED 3X confocal microscope. Cells were imaged typically by use of 2 laser wavelengths (488 nm for eGFP imaging and 560 nm for mCherry imaging).

### In vitro phase separation assay

The purified recombinant proteins were mixed at indicated concentration with LLPS buffer and 5% PEG8000 at 37 °C. The mixture was pipetted onto glass bottom dish and imaged by Leica TCS SP8 STED 3X confocal microscope equipped with 100 × 1.40 NA oil objectives.

### Fluorescence recovery after photobleaching (FRAP) assay

FRAP assay was conducted by Leica TCS SP8 STED 3X confocal microscopy. 488- or 568-nm laser beam was used to bleach the fluorescent protein at a region of interest (ROI), followed with collecting time-lapse images. Fluorescence intensity of indicated ROI was measured and normalized to the fluorescence intensity of pre-bleaching image by Leica AS Lite.

### m^6^A dot blots

Total RNA was extracted with TRIzol reagent. Equal amounts of RNA (300 ng) were denatured at 95 °C for 3 min. The samples were immediately chilled on ice, added to a positively charged nylon membrane (PALL), and then cross-linked with a UV crosslinker. After blocking and incubating with the m^6^A antibody (CST) overnight, the membrane was incubated with the secondary antibody at room temperature for 1 hr. Signals were detected using a chemiluminescence imaging system. Methylene blue in 0.3 M sodium acetate (pH 5.2) was used to indicate the amount of total RNA.

### RNA immunoprecipitation assay

RIP was conducted using a RIP assay Kit (MBL) following the manufacturer’s instructions. In brief, Protein A/G magnetic beads coated with 5 μg of specific antibody or normal IgG were incubated with cell lysates at 4 °C overnight. Proteins were then extracted for immunoblot analysis, and the co-precipitated RNAs were isolated by elution buffer and purified by TRIzol reagent, and subsequently subjected to qRT‒PCR analysis.

### DNA/RNA pull-down assay

Biotin-labeled DNA and RNA probes were synthesized by RiboBio. For DNA pull-down assay, Flag-tagged NF-κB p65 expression constructs were transfected into 293T cells. At 24 hrs post-transfection, whole cell lysates were extracted from 293T cells using IP lysis buffer. Biotin-coupled DNA-protein complex was pulled down by incubating whole cell lysates with high-capacity streptavidin agarose beads (Thermo Fisher) according to the manufacturer’s instructions. The bound proteins were eluted and used for immunoblot analysis.

For RNA pull-down assay, biotin-coupled RNA complex was pulled down by incubating cell lysates with high-capacity streptavidin agarose beads, bound proteins were then extracted for immunoblot analysis.

### m^6^A RNA-IP-qRT–PCR (MeRIP‒qPCR)

An MeRIP assay was performed following a previously described procedure with minor modifications (60). Briefly, 200 μg of total RNA was fragmentated to approximately 150-200 nt in length, purified by magnetic beads, and then incubated with anti-m^6^A antibody- or IgG-conjugated protein A/G magnetic beads in 1 × IP buffer at 4 °C overnight. Immunoprecipitated methylated RNAs were competitive eluted by free m^6^A, and recovered with a RNeasy kit (QIAGEN). One-tenth of the fragmented RNA was saved for use as the input control and further analyzed by qRT‒PCR with the MeRIP RNAs using primers for the targeted gene. The related enrichment of m^6^A in each sample was calculated by normalizing the number of amplification cycles (Cq) of the m^6^A-IP portion to the Cq of the corresponding input portion.

### Quantification of the m^6^A modification by LC-MS/MS

The 200 ng extracted mRNA was digested into nucleosides by Nuclease P1 (1 U, NEB, M0660S) and shrimp alkaline phosphatase (rSAP, 1 U, NEB, M0371S) in 50 μL RNase-free water at 37 °C overnight. The mixture was diluted to 100 μL, 10 μL of which was injected into an LC–MS/MS system consisting of a high-performance liquid chromatographer (Shimadzu) equipped with a C18-T column (Weltech) and a Triple Quad 4500 (AB SCIEX) mass spectrometer in positive ion mode by multiple-reaction monitoring. Mass transitions of m/z 268.0–136.0 (A), m/z 245.0–113.1 (U), m/z 244.0–112.1 (C), m/z 284.0–152.0 (G) and m/z 282.0–150.1 (m^6^A) were monitored. A concentration series of pure commercial nucleosides (MCE) was employed to generate standard curves. The concentration of nucleosides in samples were obtained by fitting signal intensity to a standard curve with certain ratios calculated subsequently.

### RNA-seq and data analysis

Whole-cell total RNA was isolated using TRIzol reagent and quantified using a NanoDrop 2000 spectrophotometer (Thermo). The cDNA library was constructed by Biomarker Technologies. Sequencing was performed on an Illumina HiSeq 2500 platform. High-quality reads were mapped to the human reference genome (hg19) or mouse reference genome (mm9) using HISAT2. DESeq, an R package, was applied for differential gene expression analysis. We filtered the differentially expressed genes based on a false discovery rate (FDR) <0.05.

### Methylated RNA immunoprecipitation sequencing (MeRIP-seq) and data analysis

Total RNA was extracted by TRIzol reagent and mRNA was isolated using a Dynabeads mRNA purification kit (Thermo Fisher). Then, cellar mRNA was fragmented using a fragmentation kit (Thermo Fisher), and subsequent m^6^A immunoprecipitation, MeRIP-seq and data analysis was carried out as previously described with minor modifications (81).

### Statistical analysis

The data are represented as the mean ± s.d. unless otherwise indicated, and two-tailed Student’s *t*-test or Mann–Whitney U test with a confidence interval of 95% were performed for all statistical analyses (survival curves were compared by the log-rank test) with GraphPad Prism 6 software. Differences between two groups were considered statistically significant when the *P* value was less than 0.05.

### Study approval

All animal experiments were performed in accordance with the NIH Guide for the Care and Use of Laboratory Animals (National Academies Press, 2011), with the approval of the Institutional Animal Care and Use Committee (IACUC) of Sun Yat-sen University (Guangzhou, China).

### Data and code availability

Publicly available datasets, which were analyzed in this study including GSE19315, GSE198326, GSE2411, GSE2638, GSE69063, GSE13887, GSE137268, GSE97779, GSE166388, GSE208303, GSE125527, GSE16879, GSE193193, GSE179874, GSE119600, GSE10616, GSE137218, GSE227851 and GSE189847 are available at the GEO database (https://www.ncbi.nlm.nih.gov/geo/). The validated IBDMDB data and relevant participants’ information are available at the IBDMDB database (http://ibdmdb.org). All data generated for this paper have been deposited at the SRA under access number: PRJNA943438. Values for all data points in graphs are reported in the Supporting Data Values file. See complete unedited blots in the supplemental material.

## Acknowledgments

This work was supported by projects from the National Natural Science Foundation of China (32200712, U23A6012 and 31970852), the Guangdong Science and Technology Department (2023B1212060028, 2023A1515010541 and 2023A1515010628), the Ministry of Science and Technology of the People’s Republic of China (2018YFD0900502), the “Decoding TCM” collaborative research project of BUCM (90010060920009), the Fundamental Research Funds for the Central Universities (23yxqntd001) and the National Key Research and Development Project (2019YFC1710104).

## Author contributions

YG, SY and AX designed the study, analyzed the data, and prepared the manuscript with input from the other authors. YG, RC, TL, JH, YC, YL, XX and GX performed the experiments collaboratively. BL, HC and GL performed the data analysis of MeRIP-seq. SY and AX conceived the study, supervised experiments, and wrote the paper. AX and SY led the project and finally approved the manuscript.

The authors have declared that no conflict of interest exists.

